# Haematopoietic stem cell numbers are not solely determined by niche availability

**DOI:** 10.1101/2023.10.28.564559

**Authors:** Shoichiro Takeishi, Tony Marchand, Wade R. Koba, Daniel K. Borger, Chunliang Xu, Chandan Guha, Aviv Bergman, Paul S. Frenette, Kira Gritsman, Ulrich Steidl

## Abstract

Haematopoietic stem cells (HSCs) reside in specialized microenvironments, also referred to as niches, and it has been widely believed that HSC numbers are determined by the niche size alone^1–5^. However, the vast excess of the number of niche cells over that of HSCs raises questions about this model. We initially established a mathematical model of niche availability and occupancy, which predicted that HSC numbers are restricted at both systemic and local levels. To address this question experimentally, we developed a femoral bone transplantation system, enabling us to increase the number of available HSC niches. We found that the addition of niches does not alter total HSC numbers in the body, regardless of whether the endogenous (host) niche is intact or defective, suggesting that HSC numbers are limited at the systemic level. Additionally, HSC numbers in transplanted wild-type femurs did not increase beyond physiological levels when HSCs were mobilized from defective endogenous niches to the periphery, indicating that HSC numbers are also constrained at the local level. Our study demonstrates that HSC numbers are not solely determined by niche availability, thereby rewriting the long-standing model for the regulation of HSC numbers.

## Main

Since Schofield proposed the niche model in the 1970s, in which HSCs expand until they occupy their niches^5^, it has long been assumed that HSC numbers are solely determined by niche availability. This idea is further supported by the observation that transplanted HSCs do not engraft unless available niche ‘space’ is emptied by conditioning, such as irradiation, chemotherapy or other methods, which damages or mobilizes endogenous HSCs^5–10^. We and others have previously identified several HSC niche components, such as perivascular mesenchymal stem cells (MSCs), which are marked by Nestin-GFP, Cxcl12-GFP, leptin receptor or CD51/CD140α expression^11–16^. It is also known that endothelial cells (ECs), characterized by the expression pattern of CD31, CD144, Sca-1 and CD62E, contribute to the HSC niche^11,17–19^. These cells produce niche factors, such as C-X-C motif chemokine ligand 12 (CXCL12) and stem cell factor (SCF, encoded by *Kitl*), which are essential for the retention and maintenance of HSCs in the BM. Genetic depletion of these cytokines or these HSC niches results in a reduction in HSC numbers in the BM^11–15,20,21^. One puzzling observation is that the number of defined niche cells is significantly greater than the number of HSCs^11,14–16^. Although these findings do not exclude the possibility that a small population of these niche cells creates unique saturable spaces, or that HSCs compete for space with their progenitors that also depend on niche factors, it is possible that HSC numbers are determined by other factors in addition to local control by niches. Therefore, we aimed to examine whether HSC numbers are solely determined by niche availability in this study.

### Bone transplantation provides additional HSC niches

To evaluate the relationship between niche availability and HSC numbers, we initially established a phenomenological mathematical model with total HSC numbers in the body as a function of available niche space (equivalent to the number of available niches) (see Methods; Fig. 1a). The model shows that as the number of available niches increases, the number of occupying HSCs initially increases almost linearly, which aligns with the concept of local, niche availability-driven restrictions on HSC numbers. However, this increase in HSC numbers saturates when the number of available niches reaches a critical level. Beyond this point, not all niches are occupied, regardless of available space. One potential reason for this saturation is that HSC numbers could also be limited at the systemic level, with total HSC numbers in the body being restricted. Given that our mathematical model is a predictive conceptual one without data fitting, different parameter settings yielded different numerical outcomes, but qualitatively identical behaviour was observed in all settings tested (Extended Data Fig. 1a).

**Fig. 1.**
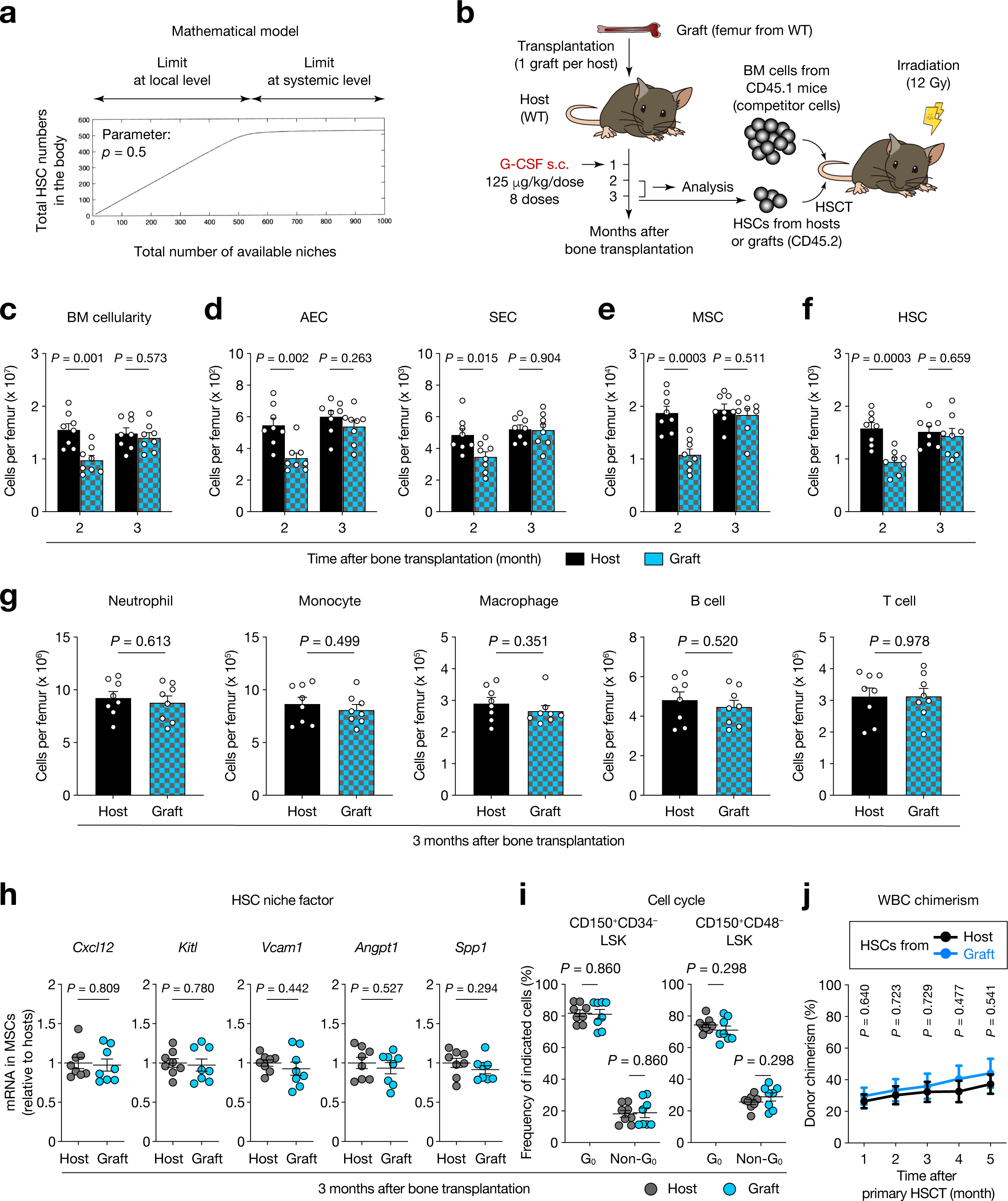
Normal functions of HSCs and their niches in the bone transplantation model. **a**, Mathematical model of niche availability and occupying HSC numbers in the body. **b**, Experimental strategy for transplantation of WT femurs to WT mice, G-CSF administration, HSC transplantation (HSCT) and analyses. **c**-**f**, The number of BM cells (**c**), ECs (**d**), MSCs (**e**) and HSCs (**f**) per host femur and graft. 8 host femurs and 8 grafts from 8 host mice at each indicated time point after bone transplantation. **g**, The number of differentiated haematopoietic cells per host femur and graft at three months after bone transplantation. 8 host femurs and 8 grafts from 8 host mice. **h**, Quantification of mRNA levels of the indicated HSC niche factors relative to *Actb* in MSCs sorted from the host femurs and the grafts at three months after bone transplantation (normalized to the host femurs). 8 host femurs and 8 grafts from 8 host mice. **i**, Frequency of quiescent (G_0_) and proliferating (non-G_0_) cells in HSCs from the host femurs and the grafts at three months after bone transplantation. 8 host femurs and 8 grafts from 8 host mice. **j**, White blood cell (WBC) chimerism (CD45.2) in recipient mice transplanted with HSCs (CD45.2) derived from the host femurs or the grafts mixed with competitor BM cells (CD45.1) at the indicated time points after primary HSCT. *n* = 10 mice per group. Data are represented as mean ± SEM. Significance was assessed using a two-tailed unpaired Student’s *t*-test.

To experimentally test this hypothesis and model, we aimed to augment the overall availability of niches *in vivo* and to assess the impact of such a gain of niches on HSC numbers. To achieve this goal, we utilized a bone transplantation method our laboratory has recently established, transplanting femoral bones from one adult mouse to another^22,23^. In this model, when femurs (hereafter referred to as grafts) from wild-type (WT) mice are transplanted into nonconditioned WT mice (called hosts), MSCs (marked by CD45^−^Ter-119^−^CD31^−^CD51^+^CD140α^+^)^16^ persist in the grafts, while phenotypic HSCs (defined by Lin^−^Sca-1^+^c-Kit^+^CD150^+^CD48^−^CD34^−^)^24^, as well as differentiated haematopoietic cells, are no longer detected within three days after transplantation (Extended Data Fig. 2a-f, Extended Data Fig. 3a-e). We also transplanted femurs from Nestin-GFP transgenic mice^14^ into Nestin-GFP mice (one graft per host) and the mice were followed up to five months after transplantation (Extended Data Fig. 4a). We confirmed the overlap of Nestin-GFP^+^ cells and CD51^+^CD140α^+^ cells in the CD45^−^Ter-119^−^CD31^−^ fraction of the grafts, as known for endogenous BM^16^ (Extended Data Fig. 4b). Based on imaging, Nestin-GFP^+^ cells were observed in the grafts as well as in the host femurs at one and five months after transplantation, with robust vascularization shown by *in vivo* staining of the BM with CD31/CD144 (Extended Data Fig. 4c). Consistent with our recent study^22^, flow cytometric analyses revealed progressive recovery of whole BM cells, MSCs and HSCs in the grafts (Extended Data Fig. 4d-f). However, while the host femurs and the grafts harboured comparable numbers of BM cells, MSCs and differentiated haematopoietic cells at five months after transplantation (Extended Data Fig. 4g), HSC numbers in the grafts were still lower than those in the host femurs.

We next examined the origin of MSCs, ECs and haematopoietic cells in the grafts at five months after transplantation. When femurs from Nestin-GFP mice were transplanted into WT mice, most CD51^+^CD140α^+^ cells in the CD45^−^Ter-119^−^CD31^−^ fraction of the grafts were positive for Nestin-GFP (Extended Data Fig. 5a, b). In contrast, Nestin-GFP^+^ cells were hardly detected in the CD51^+^CD140α^+^ fraction of WT grafts transplanted to Nestin-GFP mice (Extended Data Fig. 5c, d), indicating that Nestin-GFP^+^ MSCs in the grafts originated from the grafts. Similar experiments were then performed using Cdh5 (VE-Cadherin)-CreER; iTdTomato mice. Imaging and flow cytometric analyses confirmed TdTomato fluorescence in both arterial (CD45^−^Ter-119^−^CD31^+^Sca-1^high^CD62E^low^) and sinusoidal (CD45^−^Ter-119^−^CD31^+^Sca-1^low^CD62E^high^) ECs^19^ (AECs and SECs, respectively) regardless of whether Cdh5-CreER; iTdTomato mice were used as hosts or grafts (Extended Data Fig. 5e-j), showing that these cells were derived from both the hosts and the grafts. Next, femurs from CD45.1 mice were transplanted into CD45.2 animals, and we observed that almost all BM cells, including HSCs, expressed CD45.2 (Extended Data Fig. 5k, l), indicating that haematopoietic cells in the grafts were colonized from the hosts, consistent with our recent study^22^.

Because granulocyte colony-stimulating factor (G-CSF) mobilizes HSCs from the BM to the periphery^25^, and HSCs in grafts are of host origin, we examined the effects of administration of G-CSF on the recovery of HSCs in grafts (Extended Data Fig. 6a). Imaging analyses at three months after transplantation revealed normal BM vascular architecture in the grafted femurs, as evidenced by comparable vascular density and arteriole lengths between the host and grafted femurs (Extended Data Fig. 6b-d). We did not observe any differences in Nestin-GFP^+^ density in imaging (Extended Data Fig. 6e), which was further confirmed by flow cytometric analyses, demonstrating equivalent BM cellularity and frequency of Nestin-GFP^+^ cells in host versus grafted bones (Extended Data Fig. 6f, g). Similar observations were made when we performed the same experiments with WT hosts and grafts (Fig. 1b). BM cellularity and the number of ECs, MSCs, HSCs and differentiated haematopoietic cells were again comparable between the host and grafted femurs three months after transplantation (Fig. 1c-g). In addition, sorted MSCs from the hosts and the grafts at three months after transplantation expressed equivalent mRNA levels of canonical niche factors, such as *Cxcl12*, *Kitl*, *Vcam1*, *Angpt1* and *Spp1*, and protein levels of CXCL12 and SCF in BM extracellular fluid (BMEF) also did not differ between the host femurs and the grafts (Fig. 1h, Extended Data Fig. 7a). Consistent with normal niche function in the grafts, HSCs from these two bones exhibited similar mean fluorescence intensity (MFI) of c-Kit and CD150, as well as cell cycle status and expression levels of cell cycle regulators, which were shown to be associated with HSC functions^26–31^ (Fig. 1i, Extended Data Fig. 7b, c). BM reconstitution assays using HSCs harvested from either host or graft femurs at three months after bone transplantation showed comparable donor chimerism in the peripheral blood and BM, which was maintained upon secondary BM transplantation (BMT) (Fig. 1j, Extended Data Fig. 7d-h). Collectively, these results demonstrate that this femur transplantation system provides adult mice with additional niches, in which host-derived HSCs are able to engraft and exert their normal function.

### HSC number restriction at the systemic level

Iron distribution experiments estimate that one femur contains only 6-7% of the total BM in mice^32^ and we observed that two femurs harbour 16.6 ± 0.880% of HSCs in the total body (Extended Data Fig. 8a). Therefore, to assess how an increase in the HSC niches affects total HSC numbers in the body, six WT femurs were transplanted per WT mouse (Fig. 2a, Extended Data Fig. 8b). A sham operation was performed as a control, followed by G-CSF administration to both groups, and HSC number determination at one and three months after transplantation, respectively. The mRNA expression of inflammatory cytokines in peripheral blood cells was not elevated at three months after transplantation (Fig. 2b). BM cellularity, EC and MSC numbers, and the levels of niche factors were comparable among the femurs from sham-operated mice, and the host femurs and the grafts from bone transplantation hosts (Fig. 2c-e, Extended Data Fig. 8c-g). These data indicated that our six-femur transplantation technique enables us to add substantial HSC niches without inducing chronic systemic inflammation. HSC numbers in the grafts were also independent of their transplanted sites, and strikingly, we found that the frequency and the absolute number of phenotypic HSCs per host femur and graft femur were lower than those per femur from the sham-operated mice (Fig. 2f, g, Extended Data Fig. 8h). A BM reconstitution assay demonstrated lower donor chimerism when BM cells from the host femurs or the graft femurs from bone transplantation hosts were transplanted (Fig. 2h). To determine whether such decreased repopulation activity was attributable to lower HSC numbers or a decreased engraftment capacity of HSCs, we performed competitive repopulation experiments with sorted HSCs. We found no difference in the repopulation activities among the three groups, consistent with equivalent MFI of c-Kit and CD150 in transplanted HSCs (Fig. 2i, Extended Data Fig. 8i, j). These results suggest that the decreased BM repopulation activity in the host femurs and the graft femurs from bone transplantation hosts did not result from a decline in HSC competitiveness, but from a lower number of functional HSCs. Similarly, total HSC numbers in the entire body of bone transplantation hosts (not including those in the grafts) were lower than those in the sham-operated mice (Fig. 2j). Importantly, the sum of HSC numbers in the bone transplantation hosts and the grafts was equivalent to that in the sham-operated group (Fig. 2k). These results indicate that total HSC numbers in the body are not determined by niche availability alone, and suggest that their numbers are restricted at the systemic level (Extended Data Fig. 8k).

**Fig. 2.**
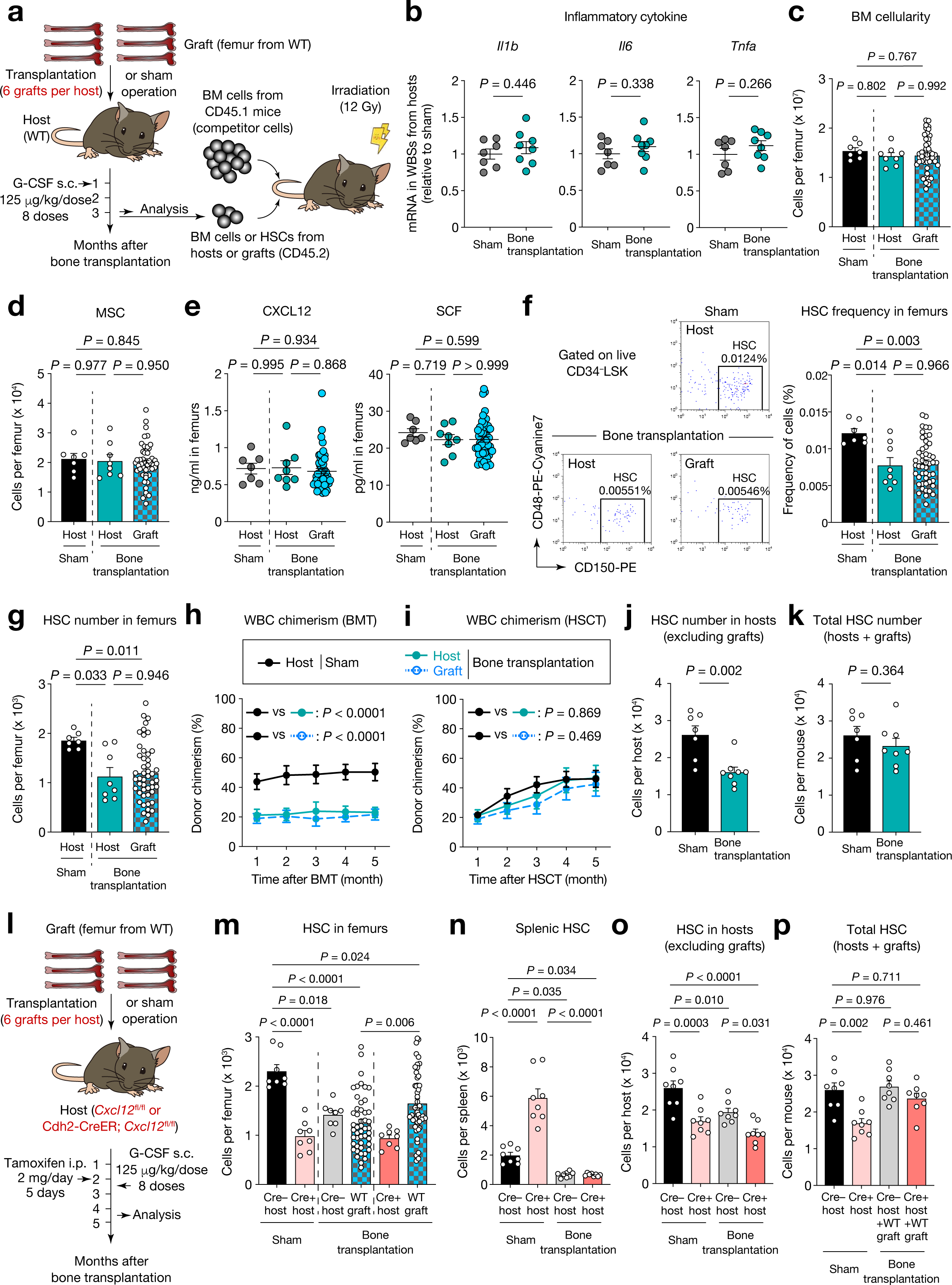
Total HSC numbers in the body remain unchanged after the niche size is increased. **a**, Schematic of transplantation of six WT femurs to WT mice and analyses. **b**, Quantification of mRNA levels of the indicated inflammatory cytokines relative to *Actb* in WBCs from sham-operated mice and bone transplantation hosts at three months after sham operation or bone transplantation (normalized to sham-operated mice). *n* = 7, 8 mice, respectively. **c**, **d**, The number of BM cells (**c**) and MSCs (**d**) per host femur and graft at three months after sham operation or bone transplantation. 7 femurs from 7 sham-operated mice, 8 host femurs and 48 grafts from 8 bone transplantation hosts. **e**, CXCL12 and SCF levels in BMEF of the host femurs and the grafts measured by ELISA at three months after sham operation or bone transplantation. 7 femurs from 7 sham-operated mice, 8 host femurs and 48 grafts from 8 bone transplantation hosts. **f**, Left: representative flow cytometry plots of HSCs isolated from the femurs of sham-operated mice (top right), host femurs (bottom left) and grafts (bottom right) of bone transplantation hosts at three months after sham operation or bone transplantation. Gates and percentages represent the frequency of the HSC population. Right: the frequency of HSCs in the host femurs and the grafts. 7 femurs from 7 sham-operated mice, 8 host femurs and 48 grafts from 8 bone transplantation hosts. **g**, Absolute HSC numbers per host femur and graft at three months after sham operation or bone transplantation. 7 femurs from 7 sham-operated mice, 8 host femurs and 48 grafts from 8 bone transplantation hosts. **h**, **i**, WBC chimerism (CD45.2) in recipient mice transplanted with BM cells (**h**) or HSCs (**i**) (CD45.2) derived from the indicated bones mixed with competitor BM cells (CD45.1). *n* = 10 mice per group. **j, k**, Absolute HSC numbers in the entire body of hosts (not including grafts) (**j**), and a sum of them in the hosts and the grafts (**k**) at three months after sham operation or bone transplantation. *n* = 7, 8 mice, respectively. **l**, Experimental strategy for transplantation of six WT femurs to Cdh2-CreER; *Cxcl12*^fl/fl^ mice and analyses. **m**, The number of HSCs per host femur and graft of the indicated genotypes in the experiment shown in **l**. 8 femurs from 8 sham-operated mice, 8 host femurs and 48 grafts from 8 bone transplantation hosts in both *Cxcl12*^fl/fl^ and Cdh2-CreER; *Cxcl12*^fl/fl^ groups. **n**-**p**, Absolute HSC numbers in the spleens (**n**), in the entire body of hosts (**o**) and a sum of them in the hosts and the grafts (**p**) of the indicated genotypes in the experiment shown in **l**. *n* = 8 mice per group. Data are represented as mean ± SEM. Significance was assessed using a two-tailed unpaired Student’s *t*-test (**b**, **j**, **k**) or one-way ANOVA (**c**-**i**, **m**-**p**).

To further investigate whether HSC numbers are constrained systemically, we next sought to determine whether total HSC numbers in the body would be maintained in hosts that harbour both defective and intact niches, taking advantage of the bone transplantation system to differentially manipulate the HSC niches in hosts and grafts. To this end, we used Cdh2 (N-Cadherin)-CreER; *Cxcl12*^fl/fl^ mice as hosts, given the known role of CXCL12 in the retention of HSCs in the BM^12^. Consistent with a study showing that Cdh2-expressing BM stromal cells contribute to the HSC niche^21^, we observed that Cdh2-CreER; TdTomato^+^ cells in the CD45^−^Ter-119^−^CD31^−^ fraction largely overlapped with CD51^+^CD140α^+^ cells and Nestin-GFP^+^ cells (Extended Data Fig. 9a, b). HSC numbers were decreased in the BM and increased in the blood and spleens of Cdh2-CreER; *Cxcl12*^fl/fl^ mice (Extended Data Fig. 9c-e), indicative of extramedullary haematopoiesis (EMH), as previously reported for *Cxcl12* flox lines that harbour Prx1-Cre or Ng2-Cre^12,20^. Based on this observation, six WT femurs were transplanted into *Cxcl12*^fl/fl^ or Cdh2-CreER; *Cxcl12*^fl/fl^ mice prior to *Cxcl12* depletion (Fig. 2l). A sham operation was performed in these genotypes in the other cohorts, and all groups were subjected to tamoxifen injection (to delete *Cxcl12* flox alleles) and G-CSF administration (to promote HSC recovery in the grafts). In the bone transplantation groups, HSC numbers per graft in Cdh2-CreER; *Cxcl12*^fl/fl^ (Cre+) hosts were greater than those in *Cxcl12*^fl/fl^ (Cre–) hosts (Fig. 2m). This could be attributable to more mobilized HSCs to the periphery after *Cxcl12* depletion in the hosts (Extended Data Fig. 9d). Notably, HSC numbers per graft in Cdh2-CreER; *Cxcl12*^fl/fl^ hosts are still lower than those in the femurs of sham-operated *Cxcl12*^fl/fl^ mice, indicating that local HSC numbers in the former bones are below the physiological level. We also observed that bone transplantation mitigated EMH in the spleens imposed by CXCL12 deficiency (Fig. 2n). In the sham-operated groups, Cdh2-CreER; *Cxcl12*^fl/fl^ mice harboured fewer total HSCs than *Cxcl12*^fl/fl^ mice (Fig. 2o), which could be attributable, at least in part, to the insufficiency of EMH in the spleens to compensate for the reduced HSC numbers in the BM. Remarkably, the sum of HSC numbers in Cdh2-CreER; *Cxcl12*^fl/fl^ hosts and WT grafts was equivalent to that in sham-operated *Cxcl12*^fl/fl^ mice (Fig. 2p). Collectively, these data suggest that total HSC numbers in the body remain unchanged when the size of the intact niche is increased, even in mice with defective niches, further supporting the notion that HSC numbers are restricted at the systemic level (Extended Data Fig. 9f).

### Local limit on HSC numbers

To further characterize how HSC numbers are determined at the systemic level, we next examined whether damage to HSCs in one part of the body would be offset by an increase in HSCs in other regions. To this end, we subjected mice to targeted irradiation of their four limbs to permanently damage the niches and thereby eradicate HSCs in the limb bones, and then assessed HSC numbers in the non-targeted areas (Fig. 3a). At three months after irradiation with 20 Gy, we confirmed that EC and MSC numbers in the limb bones were reduced (Fig. 3b, c). In contrast, these cell numbers and expression levels of niche factors in the non-targeted bone (pelvis) were not affected by irradiation (Fig. 3d-g). Although HSCs in the limbs were also decreased as expected, we did not observe any increase in HSCs in non-targeted areas (Fig. 3h, Extended Data Fig. 10a). At least two possibilities for this finding can be envisioned: 1. HSC numbers are restricted locally (in addition to limit on total HSC numbers in the body) as also predicted by our mathematical model, and 2. Systemic sensing does not counter-regulate HSC numbers when their total numbers in the body are reduced, and these two possibilities are not mutually exclusive.

**Fig. 3.**
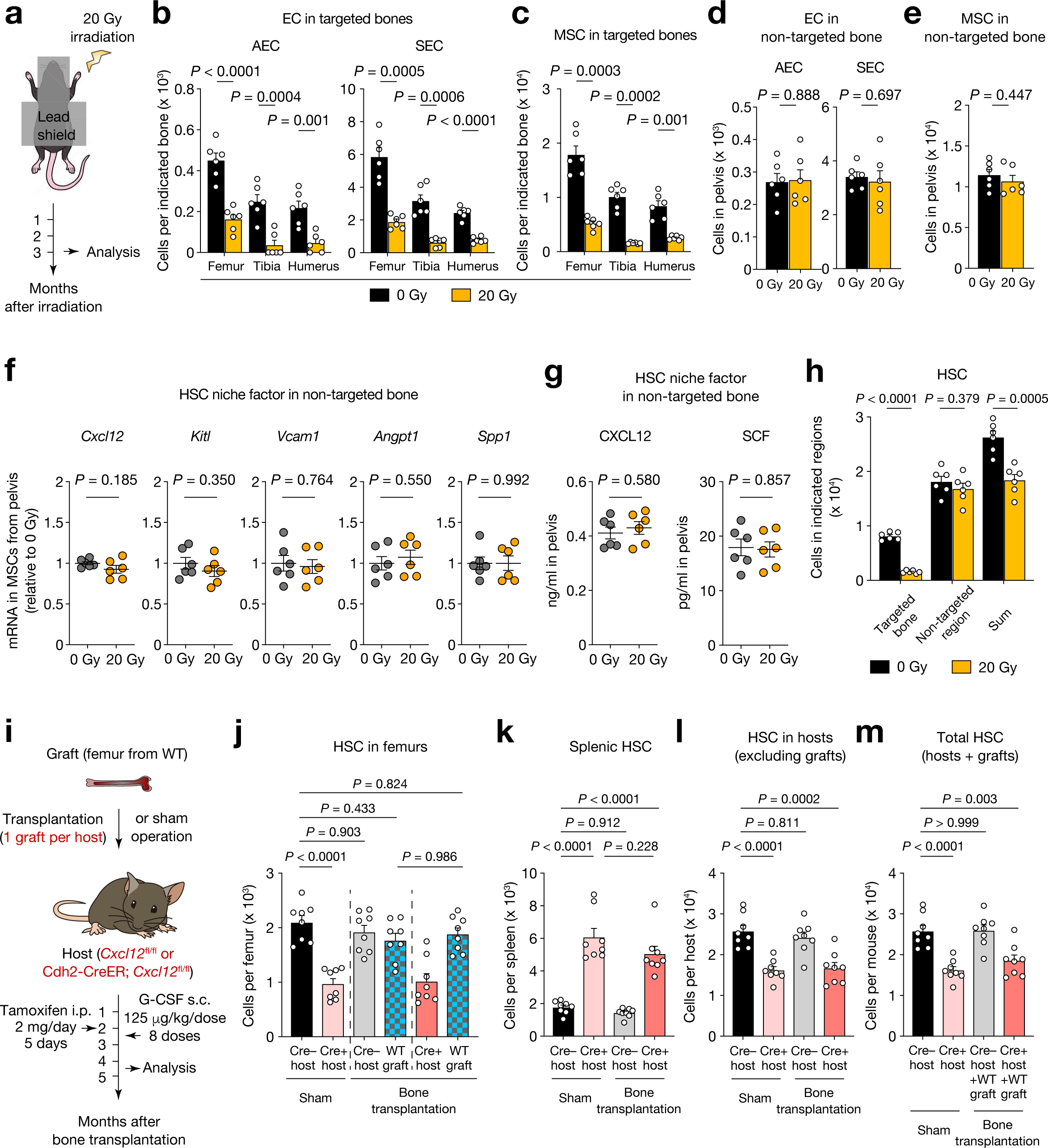
HSC numbers are limited locally. **a**, Schematic of localized irradiation to limbs at a dose of 20 Gy and analyses. **b**, **c**, The number of ECs (**b**) and MSCs (**c**) in the targeted bones (limbs) at three months after localized irradiation. *n* = 6 mice per group. **d**, **e**, The number of ECs (**d**) and MSCs (**e**) in the non-targeted bone (pelvis) at three months after localized irradiation. *n* = 6 mice per group. **f**, Quantification of mRNA levels of the indicated HSC niche factors relative to *Actb* in MSCs sorted from the non-targeted bone (pelvis) at three months after localized irradiation (normalized to 0 Gy). *n* = 6 mice per group. **g**, CXCL12 and SCF levels in BMEF of the non-targeted bone (pelvis) measured by ELISA at three months after localized irradiation. *n* = 6 mice per group. **h**, Absolute HSC numbers in the targeted bones (four limbs), non-targeted regions (skull, spine, rib cage, pelvis, spleen and liver) and a sum of them at three months after localized irradiation. *n* = 6 mice per group. **i**, Experimental strategy for transplantation of a single WT femur to Cdh2-CreER; *Cxcl12*^fl/fl^ mice and analyses. **j**, The number of HSCs per host femur and graft of the indicated genotypes in the experiment shown in **i**. 8 femurs from 8 sham-operated mice, 8 host femurs and 8 grafts from 8 bone transplantation hosts in both *Cxcl12*^fl/fl^ and Cdh2-CreER; *Cxcl12*^fl/fl^ groups. **k**-**m**, Absolute HSC numbers in the spleens (**k**), in the entire body of hosts (**l**) and a sum of them in the hosts and the grafts (**m**) of the indicated genotypes in the experiment shown in **i**. *n* = 8 mice per group. Data are represented as mean ± SEM. Significance was assessed using a two-tailed unpaired Student’s *t*-test (**b**-**h**) or one-way ANOVA (**j**-**m**).

To test the first possibility, we transplanted a single WT femur to *Cxcl12*^fl/fl^ or Cdh2-CreER; *Cxcl12*^fl/fl^ mice and compared HSC numbers per graft in these hosts (Fig. 3i), based on the assumption that the addition of only one femur would have a low impact on the total BM, and therefore HSC numbers in the grafts transplanted to *Cxcl12*^fl/fl^ mice would reach the physiological level. We reasoned that if there is no local restriction on HSC numbers, there would be more HSCs in the grafts transplanted to Cdh2-CreER; *Cxcl12*^fl/fl^ mice compared with those implanted to *Cxcl12*^fl/fl^ mice, as was seen when six femurs were transplanted to Cdh2-CreER; *Cxcl12*^fl/fl^ mice (Fig. 2m). As expected, HSC numbers per host femur and graft femur in *Cxcl12*^fl/fl^ mice were comparable to those per femur of sham-operated *Cxcl12*^fl/fl^ mice (corresponding to the physiological level) (Fig. 3j). However, CXCL12 deficiency in the host BM did not affect HSC numbers per grafted WT femur. Consistent with these data, HSC numbers in the spleens of the hosts as well as in the host body (not including the graft) were not altered by bone transplantation in the presence or absence of CXCL12 in the hosts (Fig. 3k, l). Notably, the sum of HSC numbers in the host body and the WT graft femur of Cdh2-CreER; *Cxcl12*^fl/fl^ recipient mice was lower than that of sham-operated *Cxcl12*^fl/fl^ mice (Fig. 3m), indicating that total HSC numbers in the former mice are below the systemic limit. Collectively, these data suggest that the failure to increase HSC numbers per graft in Cdh2-CreER; *Cxcl12*^fl/fl^ hosts is, at least in part, due to a local restriction on HSC numbers (Extended Data Fig. 11a).

### Systemic sensing does not compensate for HSC loss

We also tested the second hypothesis stemming from our observation in the targeted irradiation experiments, that systemic sensing may not counter-regulate HSC numbers when their total numbers in the body are reduced. To this end, we examined how HSC numbers are determined at the systemic level when local restriction on HSC numbers is lifted. We utilized Cdh2-CreER; *Kitl*^fl/fl^ mice as hosts, given that SCF is essential for the maintenance of HSCs in the BM^11,20^. We observed that tamoxifen-injected Cdh2-CreER; *Kitl*^fl/fl^ mice harboured fewer HSCs in the femurs, and equivalent numbers of haematopoietic stem and progenitor cells (HSPCs, defined by Lin^−^Sca-1^+^c-Kit^+^) in the blood and HSCs in the spleens compared with *Kitl*^fl/fl^ mice (Extended Data Fig. 12a-c), confirming that SCF from Cdh2^+^ cells indeed plays a key role in HSC maintenance in the BM. Six WT femurs were then transplanted into tamoxifen-treated *Kitl*^fl/fl^ or Cdh2-CreER; *Kitl*^fl/fl^ mice (Fig. 4a). We found that HSC numbers per graft femur in Cdh2-CreER; *Kitl*^fl/fl^ hosts were further decreased compared with those in *Kitl*^fl/fl^ hosts, and HSC numbers in the hosts exhibited a similar trend (Fig. 4b, c). Notably, the sum of HSC numbers in the hosts and the grafts in Cdh2-CreER; *Kitl*^fl/fl^ hosts was equivalent to that in sham-operated Cdh2-CreER; *Kitl*^fl/fl^ hosts, but was lower than that in sham-operated *Kitl*^fl/fl^ mice (Fig. 4d). Collectively, these data suggest that systemic sensing does not compensate for HSC loss (Extended Data Fig. 12d).

**Fig. 4.**
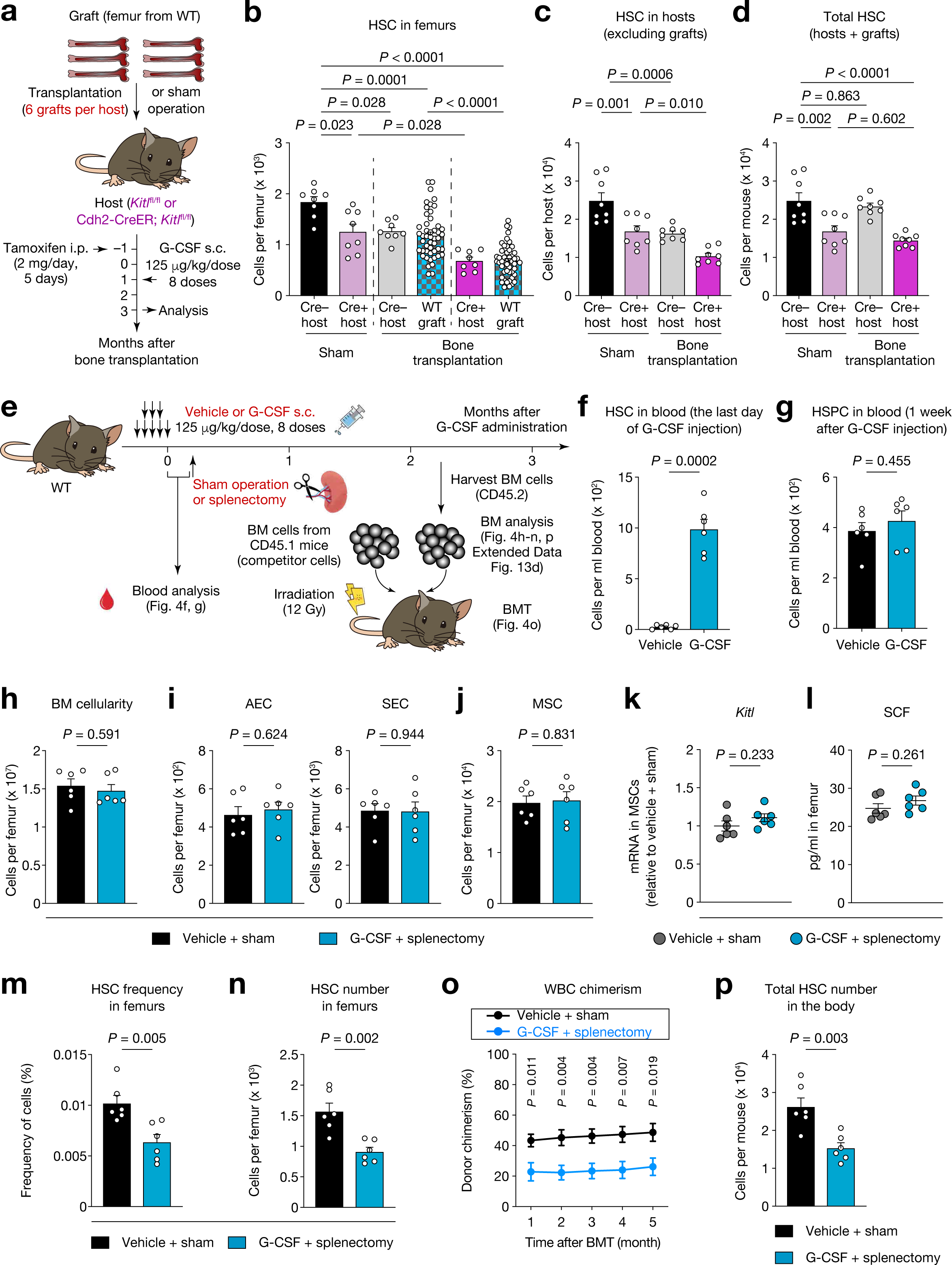
HSC number reduction is not reversed in various challenge settings. **a**, Experimental strategy for transplantation of six WT femurs to Cdh2-CreER; *Kitl*^fl/fl^ mice and analyses. **b**, The number of HSCs per host femur and graft of the indicated genotypes at three months after sham operation or bone transplantation in the experiment shown in **a**. 8 femurs from 8 sham-operated mice, 8 host femurs and 48 grafts from 8 bone transplantation hosts in both *Kitl*^fl/fl^ and Cdh2-CreER; *Kitl*^fl/fl^ groups. **c**, **d**, Absolute HSC numbers in the entire body of hosts (**c**) and a sum of them in the hosts and the grafts (**d**) of the indicated genotypes at three months after sham operation or bone transplantation in the experiment shown in **a**. *n* = 8 mice per group. **e**, Schematic of G-CSF administration, followed by splenectomy, BMT and analyses. **f**, The number of HSCs in the blood of vehicle or G-CSF-treated mice on the last day of G-CSF injection. *n* = 6 mice per group. **g**, Absolute HSPC numbers in the blood at one week after vehicle or G-CSF administration. *N* = 6 mice per group. **h**-**j**, The number of BM cells (**h**), ECs (**i**) and MSCs (**j**) per femur of the indicated cohorts at two months after G-CSF administration and splenectomy. *n* = 6 mice per group. **k**, Quantification of *Kitl* mRNA levels relative to *Actb* in MSCs sorted from the indicated cohorts at two months after G-CSF administration and splenectomy (normalized to the vehicle-treated sham-operated group). *n* = 6 mice per group. **l**, SCF levels in BMEF of the femurs from the indicated cohorts measured by ELISA at two months after G-CSF administration and splenectomy. *n* = 6 mice per group. **m**, **n**, The frequency (**m**) and the absolute number (**n**) of HSCs in the femurs of the indicated cohorts at two months after G-CSF administration and splenectomy. *n* = 6 mice per group. **o**, WBC chimerism (CD45.2) in recipient mice transplanted with BM cells (CD45.2) derived from the indicated cohorts mixed with competitor BM cells (CD45.1). *n* = 10 mice per group. **p**, Absolute HSC numbers in the entire body of the indicated cohorts at two months after G-CSF administration and splenectomy. *n* = 6 mice per group. Data are represented as mean ± SEM. Significance was assessed using a two-tailed unpaired Student’s *t*-test (**f**-**p**) or one-way ANOVA (**b**-**d**).

To exclude the possibility that rescue of HSC loss at the systemic level requires SCF, we performed splenectomy after G-CSF administration to examine the impact of removing mobilized HSCs in the spleen on the number of residual HSCs in the BM (Fig. 4e). We observed more HSCs in the blood of G-CSF-treated mice compared with vehicle-treated animals on the last day of G-CSF administration, which was no longer evident in HSPC numbers at one week after G-CSF injection (Fig. 4f, g), indicating that HSC mobilization from the BM had concluded by this point. We also confirmed that G-CSF administration decreased HSC numbers in the BM at this time point, while increasing splenic HSCs (Extended Data Fig. 13a-c), consistent with previous studies^25,33^. Subsequently, we performed splenectomy or sham operation, and analysed the BM at two months after G-CSF administration, based on a study showing that mobilized HSCs have returned to the BM two months after G-CSF treatment^25^. BM cellularity, EC and MSC numbers and the levels of niche factors, including SCF, were comparable between the vehicle + sham operation and the G-CSF + splenectomy groups (Fig. 4h-l, Extended Data Fig. 13d). Strikingly, we found that splenectomy following G-CSF treatment decreased the frequency and the absolute number of HSCs in the femurs, which was further confirmed by competitive repopulation assays of BM cells (Fig. 4m-o). Similarly, the G-CSF + splenectomy group harboured a lower number of total HSCs in the body than the vehicle + sham operation group (Fig. 4p). These results demonstrate that systemic sensing does not counter-regulate HSC numbers upon their reduction even when SCF is present in this setting. These findings suggest that such dynamics of total HSC numbers could also contribute to their reduction after targeted irradiation (Fig. 3h). Furthermore, these observations imply that the increase in HSC numbers in the 6-femur grafts due to CXCL12 deficiency in the hosts (Fig. 2m) are attributable to HSC migration from the hosts to the grafts rather than HSC expansion. Taken together, our experimental data demonstrate that HSC numbers are not solely determined by niche availability, as predicted by our mathematical model.

## Discussion

In this study, we initially established a mathematical model that predicted that the number of HSCs is determined at both the systemic and local levels. We have validated this prediction experimentally by investigating the effect of increased niche size on HSC numbers utilizing a femoral bone transplantation technique (Extended Data Fig. 14). Further investigation will be required to elucidate the molecular mechanisms underlying systemic sensing and counter-regulation of HSC numbers. In this context, it would be worthwhile to explore whether factors from outside the BM that influence HSC behaviour^34–39^ are involved in this process, given that it remains unknown whether such long-range signals maintain HSCs systemically and/or locally. Our findings appear to be inconsistent with a study reporting that transplantation of a very large number of HSPC (approximately four times the total HSC numbers in the body) without myeloablation led to an increase in HSC numbers in the recipients^40^. However, given the accumulating evidence that HSCs fail to engraft when a smaller number of them are transplanted to nonconditioned mice^5–10^, it is possible that transplantation of a hyper-physiological number of HSPCs disrupts the restriction on total HSC numbers in the body.

We also found that systemic sensing does not compensate for the reduction of total HSC numbers in the body in various settings, including in the bone transplantation system using SCF-deficient mice, unlike in stress haematopoiesis. In contrast to the disappearance of nearly all haematopoietic cells in the body after 5-FU challenge or BMT, BM cells persist in the body under the conditions tested in this study. Such a presence (in the case of localized irradiation, bone transplantation using SCF-deficient mice, or G-CSF administration + splenectomy) or absence (after 5-FU challenge or BMT) of residual BM cells in the body might contribute to the different responses to HSC reduction in these settings. Our findings indicate that HSCs are under stringent control by at least two modes of extrinsic regulation, which limit their ability to increase total HSC numbers. Given recent notions that the majority of HSCs are dispensable for steady-state haematopoiesis^41–43^ and that HSCs are involved in the pathogenesis of haematologic malignancies as a source of (pre)leukemic stem cells^44,45^, it is intriguing to speculate that the haematopoietic system might be equipped with extrinsic safeguards that suppress HSC numbers to prevent leukemogenesis, even at a slight expense of normal haematopoiesis. It has been reported that HSC niche cell numbers are increased in myeloid malignancies^46,47^. In future studies, it may be interesting to determine whether the mechanisms of HSC number regulation are affected in these or other pathophysiological conditions.

## Methods

### Mice

B6.Cg-*Gt(ROSA)26Sor^tm^*^14^(CAG–tdTomato)*^/Hze^*/J (iTdTomato) (#007914), C57BL/6J (CD45.2) (#000664) and B6.SJL-*Ptprc^a^ Pepc^b^*/BoyJ (CD45.1) (#002014) mice were purchased from The Jackson Laboratory. Nestin-GFP mice^48^ were bred in our facility. Cdh5-CreER, Cdh2-CreER, *Cxcl12*^fl/fl^, *Kitl*^fl/fl^ mice were kindly provided by R. H. Adams (Max Planck Institute for Molecular Biomedicine, Germany), L. Li (Stowers Institute for Medical Research, USA), T. Nagasawa (Osaka University, Japan) and S. J. Morrison (University of Texas Southwestern Medical Center, USA), respectively. Unless indicated otherwise, 8–10-week-old mice of both genders were used for experiments. All these mice were backcrossed with C57BL/6J mice for more than 10 generations and maintained in pathogen-free conditions under a 12 h/12 h light/dark cycle, at a temperature of 21 ± 1°C and humidity of 40–70%, and were fed with autoclaved food and water. This study complied with all ethical regulations involving experiments with mice, and all experimental procedures performed on mice were approved by the Institutional Animal Care and Use Committee of Albert Einstein College of Medicine. No randomization or blinding was used to allocate experimental groups.

### Femoral bone transplantation

Femurs with intact periosteum were isolated from 8–10-week-old donor mice and preserved on ice in phosphate-buffered saline (PBS; 21-040-CV, Corning) until they were implanted in recipient mice. For transplantation of a single femur, nonconditioned recipient mice that were age and sex-matched with donor mice were anaesthetized with ketamine and xylazine, and a small incision was made at their unilateral thoracic region. Subsequently, the preserved femur was implanted subcutaneously, and the wound was closed. For the transplantation of six femurs, small incisions were made at the bilateral thoracic, inguinal and neck regions of recipient mice, and then one femur was implanted in each area, followed by the wound closure. A sham operation was performed by making small incisions on the same area of skin as the control bone transplantation group and closing them.

### Splenectomy

After mice were anaesthetized with ketamine and xylazine, a longitudinal incision was made in the skin and peritoneum on the left dorsolateral side of the abdomen, caudal to the last rib. The splenic artery was ligated and the spleen was removed. The abdominal wall was then closed and the skin was sutured. A sham operation was performed by exteriorizing the spleen and then reinserting it into the abdominal cavity.

### *In vivo* treatment

For G-CSF treatment, G-CSF (NEUPOGEN/Filgrastim; 300 µg/ml, purchased from Jack D. Weiler Hospital of Albert Einstein College of Medicine) was injected subcutaneously (s.c.) at a dose of 125 μg kg^−1^ twice a day (8 divided doses) beginning in the evening of the first day. When used in bone transplantation experiments, G-CSF was administered to all groups at one month after femurs were implanted or a sham operation was performed otherwise indicated. When HSC mobilization was checked, blood was collected at 3 h or 7 days after the final morning dose. For induction of CreER-mediated recombination, 8-10-week-old Cdh5-CreER; iTdTomato mice were injected intraperitoneally (i.p.) with 2 mg tamoxifen (T5648, Sigma-Aldrich) dissolved in corn oil (C8267, Sigma-Aldrich) for five consecutive days (10 mg in total per mouse). Four weeks after the injection, these mice were used as hosts, or femurs were isolated for transplantation. In experiments examining the overlap of Cdh2^+^ cells and MSCs, 8-10-week-old Cdh2-CreER; iTdTomato or Cdh2-CreER; iTdTomato; Nestin-GFP mice were injected with tamoxifen and subjected to analyses four weeks after the injection. In experiments using Cdh2-CreER; *Cxcl12*^fl/fl^ or *Cxcl12*^fl/fl^ mice as hosts, tamoxifen was administered at two months after femurs were implanted or a sham operation was performed to these mice. In experiments using Cdh2-CreER; *Kitl*^fl/fl^ or *Kitl*^fl/fl^ mice as hosts, tamoxifen was administered to four to five-week-old mice before femurs were implanted or a sham operation was performed.

### Whole-mount imaging of host femurs and femoral grafts

Antibodies used for immunofluorescence staining of femoral grafts and host femurs are CD31 (PECAM1)-Alexa Fluor 647 (MEC13.3; 102516) and CD144 (Cdh5)-Alexa Fluor 647 (BV13; 138006) from BioLegend. For all imaging experiments, these antibodies (5 μg each) were injected to mice via the retro-orbital plexus for the vasculature staining, and mice were euthanized 10 min after injection. Femoral grafts and host femurs were then isolated and fixed in 4% paraformaldehyde (PFA; 15710, Electron Microscopy Sciences) overnight at 4°C. For cryopreservation, the bones were incubated sequentially in 10%, 20% and 30% sucrose/PBS at 4°C for 1 h each, embedded and flash frozen in SCEM embedding medium (C-EM002, SECTION-LAB Co. Ltd.), and stocked at –80°C. For whole-mount imaging, bones were placed at –20°C overnight, and shaved with a Cryostat (CM3050, Leica) until the BM cavity was fully exposed. Sections were carefully harvested from the melting embedding medium, rinsed with PBS, and postfixed with 4% cold PFA for 10 min followed by permeabilization in 0.5% Triton X-100/PBS for 3 h at room temperature (20–25°C) and incubation with 2 µg ml^-1^ 4′,6-diamino-2-phenylindole (DAPI; D9542, Sigma-Aldrich) for 30 min. Images were acquired at room temperature using a Zeiss Axio examiner D1 microscope (Zeiss) with a confocal scanner unit (Yokogawa), and reconstructed in three dimensions with SlideBook 6 software (Intelligent Imaging Innovations) and the Fiji build of ImageJ (National Institute of Health, NIH).

### Cell preparation

For analyses of haematopoietic cells in host femurs and femoral grafts, BM cells in these bones were flushed and dissociated using a 1-ml syringe with PBS via a 21-gauge needle. For analyses of haematopoietic cells throughout the mouse body, BM cells in femurs, tibias, humeri and pelvis were harvested by flushing and dissociating, and radii, skull, spine, sternum and ribs were minced into small pieces with scissors, crushed with a mortar and pestle, and filtered through a 70 µm cell strainer. Splenic cells were obtained by gentle grinding with slide glasses and passing through a 70 µm cell strainer. Cells in the liver were obtained by gentle grinding with slide glasses followed by digestion at 37°C for 30 min in 1 mg ml^−1^ collagenase type IV (17104019, Gibco), 2 mg ml^−1^ dispase (17105041, Gibco) and 50 μg ml^−1^ DNase I (DN25, Sigma-Aldrich). Peripheral blood was collected by retro-orbital bleeding of mice anaesthetized with isoflurane, and mixed with EDTA to prevent clotting. The data from the bones above, spleen, liver and blood (assumed to be 2 ml per animal) were summed to calculate the total HSC numbers in the mouse body. For analyses of BM stromal cells, intact flushed BM plugs were digested at 37°C for 30 min in 1 mg ml^−1^ collagenase type IV, 2 mg ml^−1^ dispase and 50 μg ml^−1^ DNase I in Hank’s balanced salt solution (HBSS) with calcium and magnesium (21-023-CV, Gibco). These single-cell suspensions were then subjected to red blood cell lysis with ammonium chloride and washed in ice-cold PEB (PBS containing 0.5% BSA and 2 mM EDTA).

### Flow cytometric analysis and cell sorting

Cells were surface-stained in PEB for 30-60 min at 4°C. Antibodies used for flow cytometric analyses and sorting are anti-CD45-APC-eFluor 780 (30-F11; 47-0451-82), anti-Ter-119-APC-eFluor 780 (TER-119; 45-5921-82), anti-CD31-PE-Cyanine7 (390; 25-0311-82), anti-CD51-biotin (RMV-7; 13-0512-85), anti-CD140a (PDGFRA)-PE (APA5; 12-1401-81), anti-CD140a-PE-Cyanine7 (APA5; 25-1401-81), anti-Ly6A/E (Sca-1)-FITC (D7; 11-5981-82), anti-Ly-6G/Ly-6C (Gr-1)-FITC (RB6-8C5; 11-5931-85), anti-Ly-6G/Ly-6C-APC-eFluor 780 (RB6-8C5; 47-5931-82), anti-CD11b-PE (M1/70; 12-0112-83), anti-CD11b-PE-Cyanine7 (M1/70; 25-0112-82), anti-CD11b-APC-eFluor 780 (M1/70; 47-0112-82), anti-B220-APC-eFluor 780 (RA3-6B2; 47-0452-82), anti-CD3e-APC-eFluor 780 (145-2C11; 47-0031-82), anti-CD48-PerCP-eFluor 710 (HM48-1; 46-0481-85), anti-CD48-PE-Cyanine7 (HM48-1; 25-0481-80), anti-CD34-eFluor 660 (RAM34; 50-0341-82; 1:50 dilution), anti-CD115-APC (AFS98; 17-1152-82) and anti-CD45.1-PE-Cyanine7 (A20; 25-0453-82) from eBioscience, anti-CD62E-PE (10E9.6; 553751) from BD Biosciences, anti-CD117 (c-Kit)-PE-Cyanine7 (2B8; 105814), anti-CD117-Brilliant Violet 421 (2B8; 105828), anti-CD150-PE (TC15-12F12.2; 115904), F4/80-PE (BM8; 123110) and anti-CD45.2-APC (104; 109814) from BioLegend, and anti-CD3e-PerCP-Cyanine5.5 (145-2C11; 65-0031-U100) from Tonbo Biosciences. Streptavidin FITC (11-4317-87) and Streptavidin PerCP-eFluor 710 (46-4317-82) were purchased from eBioscience. Unless otherwise specified, all antibodies, Streptavidin FITC and Streptavidin PerCP-eFluor 710 were used at a 1:100 dilution. Flow cytometric analyses were carried out on BD LSRII (BD Biosciences) and cell sorting experiments were performed using BD FACSAria (BD Biosciences). Dead cells and debris were excluded by forward scatter (FSC), side scatter (SSC) and DAPI staining (1 µg ml^-1^) profiles. Data were analysed with FACS Diva 6.1 (BD Biosciences) and FlowJo (LLC) software.

### Cell cycle analysis

Single-cell suspension was stained for cell surface markers, and subsequently fixed and permeabilized with BD Cytofix/Cytoperm solution (554714, BD Biosciences) according to the manufacturer’s instructions. The cells were then stained with DAPI (Sigma-Aldrich) at 5 μg ml^−1^ and anti-Ki-67-PerCP eFluor 710 antibody (SolA15; 46-5698-80, eBioscience) or anti-Ki-67-eFluor 660 antibody (SolA15; 50-5698-82, eBioscience) at 1:100 dilution for 30 min at 4°C. After washing, the cells were analyzed in BD LSRII (BD Biosciences). A DAPI^low^Ki-67^low^ fraction was designated as the G_0_ phase of the cell cycle.

### Blood cell analysis

Peripheral blood was diluted in PBS and blood parameters were determined with Advia120 Hematology System (Siemens).

### Competitive BM and HSC transplantation

Competitive repopulation assays were performed using the CD45.1/CD45.2 congenic system. CD45.1 recipient mice were lethally irradiated (12 Gy, two split doses at least three hours apart) in a Cesium Mark 1 irradiator (JL Shepherd & Associates). For BM repopulation assays, 1 × 10^6^ CD45.2 donor-nucleated BM cells were transplanted into irradiated recipients together with 1 × 10^6^ CD45.1 BM cells. For HSC repopulating assays, 200 HSCs (CD45.2) were sorted from BM cells and transplanted into irradiated CD45.1 recipients together with CD45.1 competitor BM cells calculated to contain 200 HSCs (1:1 HSC ratio). For secondary BM transplantation, 3 × 10^6^ BM cells from primary recipient mice were transplanted into newly irradiated (12 Gy) CD45.1 recipients. CD45.1/CD45.2 chimaerism of the myeloid (CD11b^+^), B (B220^+^) and T (CD3ε^+^) lineages in recipient blood was analysed up to five months after BM or HSC transplantation using a flow cytometer, and that of BM cells was checked at five months after BM or HSC transplantation, at which the mice were euthanized.

### Targeted limb irradiation

Animals were anaesthetized by isoflurane before irradiation using the Small Animal Radiation Research Platform, SARRP (XStrahl). The orthovoltage x-ray unit operates at 220 kVp and 13 mA. Before irradiation, a static x-ray scan was acquired using 50 kVp and 0.7 mA tube current with Al filtration. Mice were maintained in a circular lucite jig with whole-body lead shielding (to protect the individualized compartments from unwanted irradiation) and ports through which secured four limbs protruded and were irradiated to 20 Gy in a single fraction.

### RNA extraction and RT and real-time PCR analysis

2,000 MSCs or HSCs were sorted directly into lysis buffer and stored at –80°C. mRNA was extracted using the Dynabeads mRNA DIRECT Purification Kit (61012, Invitrogen) according to the manufacturer’s protocols. Conventional reverse transcription with random hexanucleotide primers was then performed using the RNA to cDNA EcoDry Premix (639549, TaKaRa) in accordance with the manufacturer’s instructions. Quantitative real-time PCR was performed in 384-well plates with FastStart Universal SYBR Green Master Mix (04913914001, Roche) on QuantStudio 6 Flex Real-Time PCR System (Applied Biosystems). The PCR protocol started with one cycle at 95°C (10 min) and continued with 40 cycles at 95°C (15 s) and 60°C (1 min). All mRNA abundance was calculated relative to the corresponding amount of *Actb* (encoding β-actin) using the ΔCt method. The primer sequences are listed in Extended Data Table 1.

### ELISA

BMEF was collected by flushing the BM of one femur or pelvis using 1 ml of PBS and subsequently pelleting the cells by centrifugation. The resulting supernatant was transferred to another tube and stored at −80°C until analysis. Cytokine levels in BMEF were then measured using mouse CXCL12/SDF-1 alpha (MCX120) and SCF (MCK00) Quantikine ELISA kits (R&D Systems) according to the manufacturer’s protocols.

### Mathematical modelling of niche availability and HSC numbers

A logistic model was introduced to assess the relationship between niche availability and HSC numbers. This model aimed at bringing to light the qualitative behaviour of the dynamics of HSC niche occupancy, rather than simply experimental data fitting to a predetermined functional form. This was achieved by using the following non-linear coupled differential equations, which modelled the total number of HSCs in the body, *N*, with respect to niche space, *v*, new HSCs, *Nn*, and the production rate of HSCs, *p*.

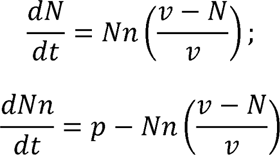

To obtain the total number of HSCs in the body as a function of the number of available niche spaces, the above equations were solved at equilibrium. The x-axis in Fig. 1a and Extended Data Fig. 1a represents available niche space (the free variable, corresponding to the number of available niches), while the y-axis shows the total number of HSCs in the body (the dependent variable).

### Statistics and reproducibility

All data are presented as mean ± SEM. n represents the number of mice in each experiment, as detailed in the figure legends, and experiments presented were successfully reproduced at least three times. No statistical method was used to predetermine sample sizes, and sample sizes were determined by previous experience with similar models of haematopoiesis, as shown in previous experiments performed in our laboratory^13,14,16,19,20,47^. Sample exclusion was only done as a result of premature mouse death. Statistical significance was determined by an unpaired, two-tailed Student’s *t*-test to compare two groups or a one-way ANOVA with Tukey’s multiple comparisons tests for multiple group comparisons. Data statistical analyses and presentation were performed using GraphPad Prism 10 (GraphPad Software), SlideBook Software 6 (Intelligent Imaging Innovations) and FlowJo (LLC).

The data in Figures 2j and 2k were obtained in the same experiments and data from the sham-operated mice were reused in each of these figure panels. The data in Figures 2o and 2p were obtained in the same experiments and data from the sham-operated *Cxcl12*^fl/fl^ and Cdh2-CreER; *Cxcl12*^fl/fl^ mice were reused in each of these figure panels. The data in Figures 3l and 3m were obtained in the same experiments and data from the sham-operated *Cxcl12*^fl/fl^ and Cdh2-CreER; *Cxcl12*^fl/fl^ mice were reused in each of these figure panels. The data in Figures 4c and 4d were obtained in the same experiments and data from the sham-operated *Kitl*^fl/fl^ and Cdh2-CreER; *Kitl*^fl/fl^ mice were reused in each of these figure panels.

### Reporting summary

Further information on research design is available in the Nature Research Reporting Summary linked to this paper.

### Data availability

Source data are provided with the paper.

## Acknowledgements

We dedicate this achievement to Dr. Paul S. Frenette, who conceived and initially supervised the study but unfortunately passed away in July 2021. We will forever be indebted to him for his brilliant mentorship and leadership. His rigorous approach and unparalleled enthusiasm for science, especially for this project aiming to answer the long-standing question in the stem cell field, laid the foundation for this manuscript. He will be deeply missed by all, and his passion for science will be carried on by his mentees and colleagues.

We are grateful to R.H. Adams, L. Li, T. Nagasawa and S. J. Morrison for providing Cdh5-CreER, Cdh2-CreER, *Cxcl12*^fl/fl^ and *Kitl*^fl/fl^ mice, respectively. We thank N. Asada, F. Nakahara, T. Mizoguchi and S. Pinho for advice with the initial experiments, and C. Prophete, P. Ciero, C. D. Cruz, A. Landeros, G. Amatuni, J. Kazmi and M. Bhuiyan for technical assistance, L. Tesfa for assistance in cell sorting, and N. P. Brodin for the targeted limb irradiation experiments. We also thank members of the Frenette laboratory, B. Will, A. Skoultchi and T. Shichita for helpful discussions and advice. This work was supported by R01 grants from the NIH (DK056638 to P.S.F. and K.G., DK112976, DK116312, HL069438 and HL116340 to P.S.F., CA253127 to U.S.), the New York State Department of Health NYSTEM IIRP (C029154 and C029570 to P.S.F.), and the Albert Einstein Cancer Center core support grant (P30CA013330). U.S. holds the Edward P. Evans Endowed Professorship in Myelodysplastic Syndromes at Albert Einstein College of Medicine. The Endowed Professorship was supported by a grant from the Edward P. Evans Foundation. This work was supported by Jane A. and Myles P. Dempsey. S.T. was supported by the Japan Society for the Promotion of Science (JSPS) Postdoctoral Fellowship for Research Abroad, the Uehara Memorial Foundation Research Fellowship and the NYSTEM Empire State Institutional Program in Stem Cell Research, and T.M. was supported by the Fondation ARC pour la Recherche sur le Cancer, the Association pour le Développement de l’Hématologie Oncologie, the Société Française d’Hématologie and the Philip foundation. D.K.B. is the recipient of a T32 NIH training grant and an F30 National Cancer Institute (NCI) predoctoral M.D./Ph.D. fellowship.

## Author contributions

S.T. conceived the project, designed the study, performed most of the experiments, analysed and interpreted the data, and wrote the manuscript. T.M., D.K.B. and C.X. helped with experiments. W.R.K. performed the targeted limb irradiation, and C.G. provided input on its experimental design. A.B. established the mathematical model of niche availability and HSC numbers, advised on the study, and edited the manuscript. P.S.F. supervised and funded the project, discussed data, and commented on the figure plan. K.G. and U.S. supervised and funded the study, analysed and interpreted data, and edited the manuscript. All authors discussed the results and reviewed the manuscript.

## Competing interests

T.M. serves as a consultant for Astellas, Jazz Pharmaceuticals, Servier and Sobi, and has received personal fees from Jazz Pharmaceuticals and Servier outside the submitted work. C. G. has received grants and personal fees from Janssen and Varian, and grants from Celldex outside the submitted work. P.S.F. served as a consultant for Pfizer, received research funding from Ironwood Pharmaceuticals outside the submitted work, and was a shareholder of Cygnal Therapeutics. K.G. has received research funding from ADC Therapeutics and iOnctura outside the submitted work. U.S. has received grants from GlaxoSmithKline, Bayer Healthcare, Aileron Therapeutics and Novartis, and personal fees from GlaxoSmithKline, Bayer Healthcare, Celgene, Aileron Therapeutics, Stelexis Therapeutics, Pieris Pharmaceuticals, Trillium and Pfizer outside the submitted work. U.S. has equity ownership in and is serving on the board of directors of Stelexis Therapeutics outside the submitted work. The rest of the authors declare no competing interests.

## Additional information

**Correspondence and requests for materials** should be addressed to Shoichiro Takeishi or Ulrich Steidl.

## Extended data figure legends

**Extended Data Fig. 1 | Mathematical model with different parameter settings exhibits qualitatively identical behaviour.**

**a**, Mathematical model of niche availability and occupying HSC numbers in the body with different parameter settings from Fig. 1a.

**Extended Data Fig. 2 | Representative flow cytometry gates used to isolate the MSCs and haematopoietic stem and differentiated cell populations.**

**a**, CD45^−^Ter-119^−^CD31^−^CD51^+^CD140α^+^ MSCs^16^.

**b**, Lineage^−^Sca-1^+^c-Kit^+^ (LSK) HSPCs and LSK CD150^+^CD48^−^CD34^−^ HSCs^24^.

**c**, Gr-1^high^CD115^low^SSC^high^ neutrophils^49^, Gr-1^low^CD115^low^F4/80^+^SSC^low^ macrophages^50^. FSC-A, forward-scatter area; SSC-A, side-scatter area.

**d**, CD115^high^CD11b^+^ monocytes^50^.

**e**, CD11b^−^B220^+^ B cells^49^.

**f**, CD11b^−^CD3ε^+^ T cells^49^.

**Extended Data Fig. 3 | While MSCs survive in the femoral grafts, virtually all haematopoietic cells are not detected shortly after bone transplantation.**

**a**, Experimental strategy to determine the number of MSCs, HSCs and differentiated haematopoietic cells in the grafts shortly after bone transplantation from WT to WT mice.

**b**, **c**, Absolute number of MSCs (**b**) and HSCs (**c**) in the grafts at the indicated time points after bone transplantation. *n* = 6, 7 grafts from 6, 7 host mice, respectively, at each time point.

**d**, Relative number of MSCs and HSCs in the grafts compared with non-transplanted femurs at the indicated time points after bone transplantation. *n* = 6, 7 grafts from 6, 7 host mice, respectively, at each time point.

**e**, The number of differentiated haematopoietic cells in the grafts at the indicated time points after bone transplantation. *n* = 8 grafts from 8 host mice at each time point.

Data are represented as mean ± SEM. Significance was assessed using a two-tailed unpaired Student’s *t*-test (**d**) or one-way ANOVA (**b**, **c**, **e**).

**Extended Data Fig. 4 | BM stroma regeneration and haematopoietic recovery in the femoral grafts.**

**a**, Schematic illustration of transplantation of Nestin-GFP femurs to Nestin-GFP mice and analyses.

**b**, Representative flow cytometry plots and the quantification of overlap of CD51^+^CD140α^+^ cells and Nestin-GFP^+^ cells in CD45^−^Ter-119^−^CD31^−^ fraction of the host femurs and the grafts at five months after bone transplantation. 8 host femurs and 8 grafts from 8 host mice. **c**, Representative confocal z-stack projection montages of Nestin-GFP (green) host femur and graft stained for double-positive CD31^+^CD144^+^ (white) vasculature with anti-CD31 and anti-CD144 antibodies at one or five months after bone transplantation. Scale bars, 1000 µm; four independent experiments yielded similar results.

**d**-**f**, The number of BM cells (**d**), MSCs (**e**) and HSCs (**f**) per host femur and graft at the indicated time points after bone transplantation. 8 host femurs and 8 grafts from 8 host mice at each time point.

**g**, The number of differentiated haematopoietic cells per host femur and graft at five months after bone transplantation. 8 host femurs and 8 grafts from 8 host mice.

Data are represented as mean ± SEM. Significance was assessed using a two-tailed unpaired Student’s *t*-test.

**Extended Data Fig. 5 | Determination of the origin of the cells in the femoral grafts.**

**a**, Experimental strategy to determine whether MSCs in the grafts are derived from the grafts. **b**, Left: representative flow cytometry plots of Nestin-GFP^+^ cells in CD45^−^Ter-119^−^CD31^−^CD51^+^CD140α^+^ MSCs isolated from the host femurs (top) and the grafts (bottom) at five months after bone transplantation in the experiment shown in **a**. Right: the frequency of the Nestin-GFP^+^ population within the CD45^−^Ter-119^−^CD31^−^CD51^+^CD140α^+^ fraction in the host femurs and the grafts. 6 host femurs and 6 grafts from 6 host mice.

**c**, Experimental strategy to determine whether MSCs in the grafts are derived from the hosts. **d**, Left: representative flow cytometry plots of Nestin-GFP^+^ cells in CD45^−^Ter-119^−^CD31^−^CD51^+^CD140α^+^ MSCs isolated from the host femurs (top) and the grafts (bottom) at five months after bone transplantation in the experiment shown in **c**. Right: the frequency of the Nestin-GFP^+^ population within the CD45^−^Ter-119^−^CD31^−^CD51^+^CD140α^+^ fraction in the host femurs and the grafts. 6 host femurs and 6 grafts from 6 host mice.

**e**, Experimental strategy to determine whether ECs in the grafts are derived from the grafts.

**f**, Confocal z-stack projection of Cdh5-CreER; iTdTomato (red) graft transplanted to WT mice and stained for double-positive CD31^+^CD144^+^ (white) vasculature with anti-CD31 and anti-CD144 antibodies at five months after bone transplantation. Scale bar, 1000 µm.

**g**, Left: representative flow cytometry plots of TdTomato^+^ cells in CD45^−^Ter-119^−^CD31^+^Sca-1^high^CD62E^low^ AEC fraction isolated from the host femurs (top left) and the grafts (bottom left) and in CD45^−^Ter-119^−^CD31^+^Sca-1^low^CD62E^high^ SEC fraction isolated from the host femurs (top right) and the grafts (bottom right) at five months after bone transplantation in the experiment shown in **e**. Right: the frequency of the TdTomato^+^ cells in AEC and SEC fractions in the host femurs and the grafts. 6 host femurs and 6 grafts from 6 host mice.

**h**, Experimental strategy to determine whether ECs in the grafts are derived from the hosts.

**i**, Confocal z-stack projection of WT graft transplanted to Cdh5-CreER; iTdTomato (red) mice and stained for double-positive CD31^+^CD144^+^ (white) vasculature with anti-CD31 and anti-CD144 antibodies at five months after bone transplantation. Scale bar, 1000 µm.

**j**, Left: representative flow cytometry plots of TdTomato^+^ cells in CD45^−^Ter-119^−^CD31^+^Sca-1^high^CD62E^low^ AEC fraction isolated from the host femurs (top left) and the grafts (bottom left) and in CD45^−^Ter-119^−^CD31^+^Sca-1^low^CD62E^high^ SEC fraction isolated from the host femurs (top right) and the grafts (bottom right) at five months after bone transplantation in the experiment shown in **h**. Right: the frequency of TdTomato^+^ cells in AEC and SEC fractions in the host femurs and the grafts. 6 host femurs and 6 grafts from 6 host mice.

**k**, Schematic illustration of the determination of the origin of haematopoietic cells in the grafts.

**l**, The frequency of CD45.1^+^ and CD45.2^+^ cells in the whole BM cells, HSCs and differentiated haematopoietic cells in the host femurs and the grafts at five months after bone transplantation in the experiment shown in **k**. 6 host femurs and 6 grafts from 6 host mice.

Data are represented as mean ± SEM.

**Extended Data Fig. 6 | Normal BM structure in G-CSF-administered femoral grafts.**

**a**, Schematic of transplantation of Nestin-GFP femurs to Nestin-GFP mice, G-CSF administration and analyses.

**b**, Representative confocal z-stack projection montages of G-CSF-administered Nestin-GFP (green) host femurs and grafts stained for double-positive CD31^+^CD144^+^ (white) vasculature with anti-CD31 and anti-CD144 antibodies at three months after bone transplantation. Green arrows indicate arterioles. Scale bars, 100 µm; four independent experiments yielded similar results.

**c**, Vasculature density in the host femurs and the grafts at three months after bone transplantation, as assessed by quantification of CD31^+^CD144^+^ double-positive vascular area divided by total BM area. *n* = 30 and 34 projections in the host femurs and the grafts, respectively; 4 host femurs and 4 grafts from 4 host mice.

**d**, Arteriolar segment length in the host femurs and the grafts at three months after bone transplantation, as assessed by quantification of the length of the Nestin-GFP^+^ signal covering CD31^+^CD144^+^ double-positive arterioles. *n* = 60 and 55 projections in the host femurs and the grafts, respectively; 4 host femurs and 4 grafts from 4 host mice.

**e**, Nestin-GFP density in the host femurs and the grafts at three months after bone transplantation, as assessed by quantification of Nestin-GFP^+^ area divided by total BM area. *N* = 28 and 31 projections in the host femurs and the grafts, respectively; 4 host femurs and 4 grafts from 4 host mice.

**f**, **g**, BM cellularity (**f**) and the frequency of Nestin-GFP^+^ MSCs (**g**) from the host femurs and the grafts at three months after bone transplantation, assessed by flow cytometry in the experiment shown in **a**. 8 host femurs and 8 grafts from 8 host mice.

Data are represented as mean ± SEM. Significance was assessed using a two-tailed unpaired Student’s *t*-test. For box plots, the box spans from the 25th to 75th percentiles and the centre line is plotted at the median. Whiskers represent the minimum to maximum range.

**Extended Data Fig. 7 | G-CSF-administered grafts harbour HSCs and their niches with normal functions.**

**a**, CXCL12 and SCF levels in BMEF of G-CSF administered host femurs and grafts measured by ELISA at three months after bone transplantation. 8 host femurs and 8 grafts from 8 host mice.

**b**, MFI of c-Kit and CD150 in HSCs from G-CSF-administered host femurs and grafts at three months after bone transplantation, as assessed by flow cytometry (normalized to the hosts). 8 host femurs and 8 grafts from 8 host mice.

**c**, Quantification of mRNA levels of the indicated cell cycle regulators relative to *Actb* in HSCs sorted from G-CSF-administered host femurs and grafts at three months after bone transplantation (normalized to the hosts). 8 host femurs and 8 grafts from 8 host mice.

**d**, Blood chimerism (CD45.2) in myeloid (CD11b^+^), B (B220^+^) and T (CD3ε^+^) cells of recipient mice transplanted with HSCs (CD45.2) derived from G-CSF-administered host femurs or grafts in competition with CD45.1^+^ BM cells at the indicated time points after primary HSCT in the experiment shown in **Fig. 1b**. *n* = 10 mice per group.

**e**, **f**, BM chimerism in whole BM, myeloid, B, T cells (**e**) and HSCs (**f**) at five months after primary HSCT. *n* = 10 mice per group.

**g**, Blood chimerism (CD45.2) in total WBC, myeloid, B and T cells of recipient mice at the indicated time points after secondary BMT of 3 × 10^6^ BM cells from primary recipients. *n* = 10 mice per group.

**h**, BM chimerism in whole BM, myeloid, B and T cells at five months after secondary BMT. *n* = 10 mice per group.

Data are represented as mean ± SEM. Significance was assessed using a two-tailed unpaired Student’s *t*-test. For box plots, the box spans from the 25th to 75th percentiles and the centre line is plotted at the median. Whiskers represent the minimum to maximum range.

**Extended Data Fig. 8 | Characterization of HSCs and their niches in the host femurs and the grafts after six femur transplantation.**

**a**, HSC distribution in the mouse BM, as assessed by flow cytometry. *n* = 11 mice.

**b**, Schematic of transplanted sites of femurs.

**c**, **d**, The number of BM cells (**c**) and ECs (**d**) in the grafts at three months after bone transplantation by their transplanted site. *n* = 16 grafts per transplanted site.

**e**, The number of ECs per host femur and graft at three months after sham operation or bone transplantation. 7 femurs from 7 sham-operated mice, 8 host femurs and 48 grafts from 8 bone transplantation hosts.

**f**, The number of MSCs in the grafts at three months after bone transplantation by their transplanted site. *n* = 16 grafts per transplanted site.

**g**, CXCL12 and SCF levels in BMEF of the grafts measured by ELISA at three months after bone transplantation by their transplanted site. *n* = 16 grafts per transplanted site.

**h**, The number of HSCs in the grafts at three months after bone transplantation by their transplanted site. *n* = 16 grafts per transplanted site.

**i**, MFI of c-Kit and CD150 in HSCs from the grafts at three months after bone transplantation by their transplanted site (normalized to the grafts transplanted to the cervix). *n* = 16 grafts per transplanted site.

**j**, MFI of c-Kit and CD150 in HSCs from the host femurs and the grafts at three months after sham operation or bone transplantation (normalized to the femurs of sham-operated mice). 7 femurs from 7 sham-operated mice, 8 host femurs and 48 grafts from 8 bone transplantation hosts.

**k**, Diagram showing the results of six WT femur transplantation to WT mice.

Data are represented as mean ± SEM. Significance was assessed using a two-tailed unpaired Student’s *t*-test. For box plots, the box spans from the 25th to 75th percentiles and the centre line is plotted at the median. Whiskers represent the minimum to maximum range.

**Extended Data Fig. 9 | Cdh2-CreER; Cxcl12fl/fl mice exhibit EMH.**

**a**, Left: representative flow cytometry plots of TdTomato^+^ cells in CD51^−^CD140α^−^ cells and CD51^+^CD140α^+^ MSCs within CD45^−^Ter-119^−^CD31^−^ fraction of Cdh2-CreER; iTdTomato mice. Right: quantification of overlap of CD51^−^CD140α^−^, CD51^+^CD140α^+^ and TdTomato^+^ cells in the CD45^−^Ter-119^−^CD31^−^ fraction of Cdh2-CreER; iTdTomato mice. *n* = 4 mice.

**b**, Left: representative flow cytometry plots of TdTomato^+^ cells in Nestin-GFP^−^ cells and Nestin-GFP^+^ MSCs within CD45^−^Ter-119^−^CD31^−^ fraction of Cdh2-CreER; iTdTomato; Nestin-GFP mice. Right: Quantification of overlap of Nestin-GFP^−^, Nestin-GFP^+^ and TdTomato^+^ cells in the CD45^−^Ter-119^−^CD31^−^ fraction of Cdh2-CreER; iTdTomato; Nestin-GFP mice. *n* = 4 mice.

**c**-**e**, The number of HSCs in the femurs (**c**), blood (**d**) and spleens (**e**) of *Cxcl12*^fl/fl^ and Cdh2-CreER; *Cxcl12*^fl/fl^ mice. *n* = 7, 9 mice, respectively.

**f**, Diagram showing the results of six WT femur transplantation to Cdh2-CreER; *Cxcl12*^fl/fl^ mice.

Data are represented as mean ± SEM. Significance was assessed using a two-tailed unpaired Student’s *t*-test.

**Extended Data Fig. 10 | HSC numbers reduced in the targeted bones after localized irradiation, while they do not alter in the non-targeted regions.**

**a**, The number of HSCs in the indicated bones and the spleens at three months after localized irradiation to limbs in the experiment shown in **Fig. 3a**. *n* = 6 mice per group.

Data are represented as mean ± SEM. Significance was assessed using a two-tailed unpaired Student’s t-test.

**Extended Data Fig. 11 | HSC numbers are restricted at the local level.**

**a**, Diagram showing the results of a single WT femur transplantation to Cdh2-CreER; *Cxcl12*^fl/fl^ mice.

**Extended Data Fig. 12 | SCF from Cdh2+ cells is essential for HSC maintenance in the BM.**

**a**, The number of HSCs per femur of *Kitl*^fl/fl^ and Cdh2-CreER; *Kitl*^fl/fl^ mice. *n* = 8 mice per group.

**b**, Absolute HSPC numbers in the blood of *Kitl*^fl/fl^ and Cdh2-CreER; *Kitl*^fl/fl^ mice. *n* = 8 mice per group.

**c**, The number of HSCs in the spleens of *Kitl*^fl/fl^ and Cdh2-CreER; *Kitl*^fl/fl^ mice. *n* = 8 mice per group.

**d**, Diagram showing the results of six WT femur transplantation to Cdh2-CreER; *Kitl*^fl/fl^ mice. Data are represented as mean ± SEM. Significance was assessed using a two-tailed unpaired Student’s *t*-test.

**Extended Data Fig. 13 | HSCs in the femurs decrease shortly after G-CSF administration, while splenic HSCs increase.**

**a**, Experimental strategy to determine HSC numbers in the femurs and the spleens shortly after G-CSF administration.

**b**-**c**, The number of HSCs per femur (**b**) and spleen (**c**) at seven days after vehicle or G-CSF administration in the experiment shown in **a**. *n* = 6 mice per group.

**d**, Quantification of mRNA levels of niche factors relative to *Actb* in MSCs sorted from the indicated cohorts at two months after G-CSF administration and splenectomy in the experiment shown in **Fig. 4e** (normalized to the vehicle-treated sham-operated group). *n* = 6 mice per group.

Data are represented as mean ± SEM. Significance was assessed using a two-tailed unpaired Student’s *t*-test.

**Extended Data Fig. 14 | Restriction of HSC numbers at both systemic and local levels.**

While it has been assumed that HSC numbers are determined by niche availability alone (left), our study demonstrates that HSC numbers are limited at both the systemic and local levels (right).

**Extended Data Table 1 | Sequence of oligonucleotides used for RT and real-time PCR analyses.**

## Notes

### Competing Interest Statement

The authors have declared no competing interest.

