## Supplemental Figure for "Haematopoietic stem cell numbers are not solely determined by niche availability"

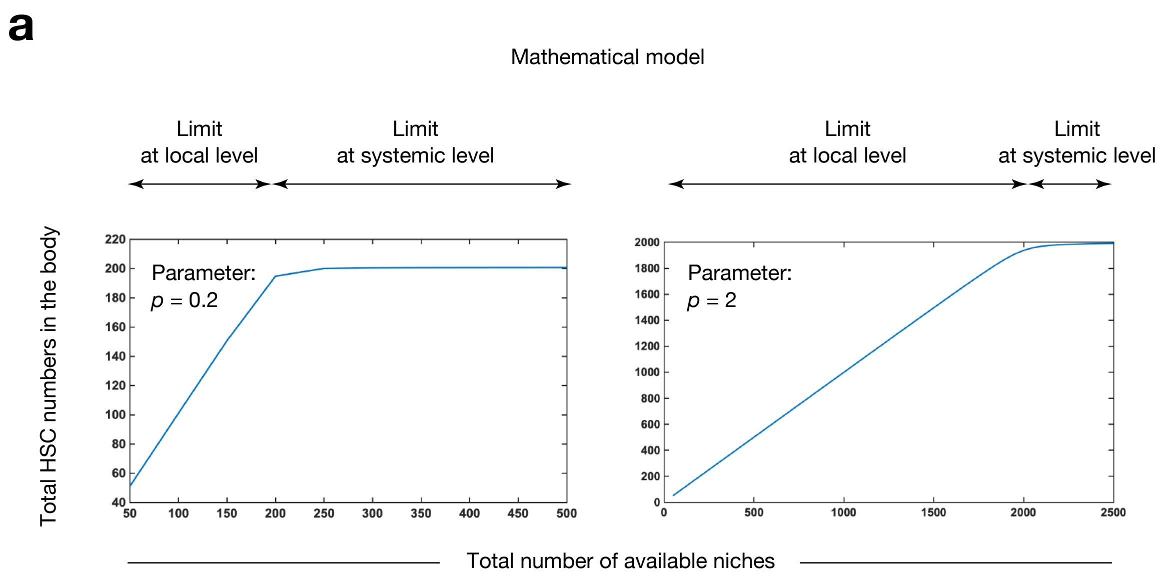


**Takeishi et al. Extended Data Fig. 1**

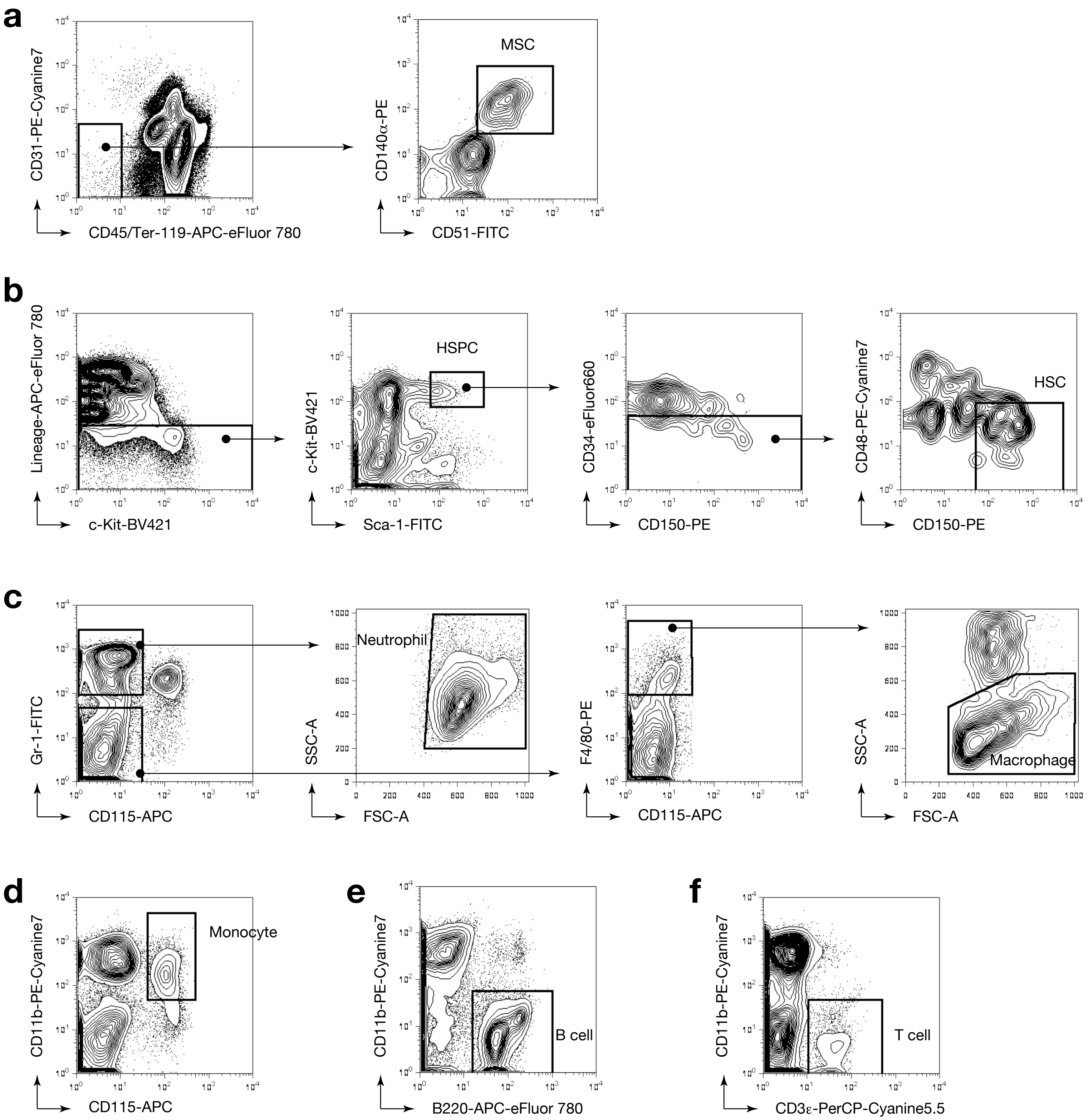


**Takeishi et al. Extended Data Fig. 2**

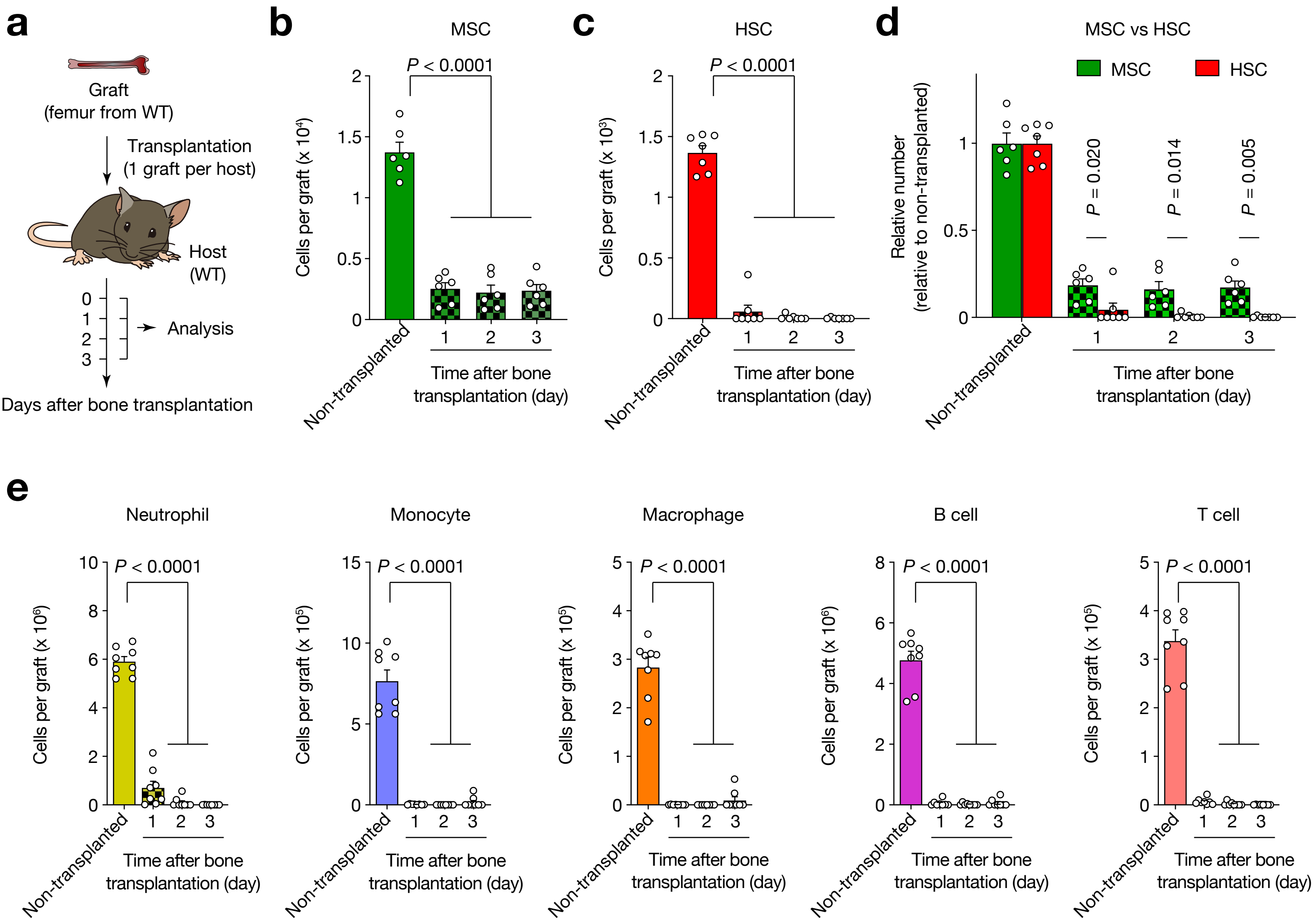


**Takeishi et al. Extended Data Fig. 3**

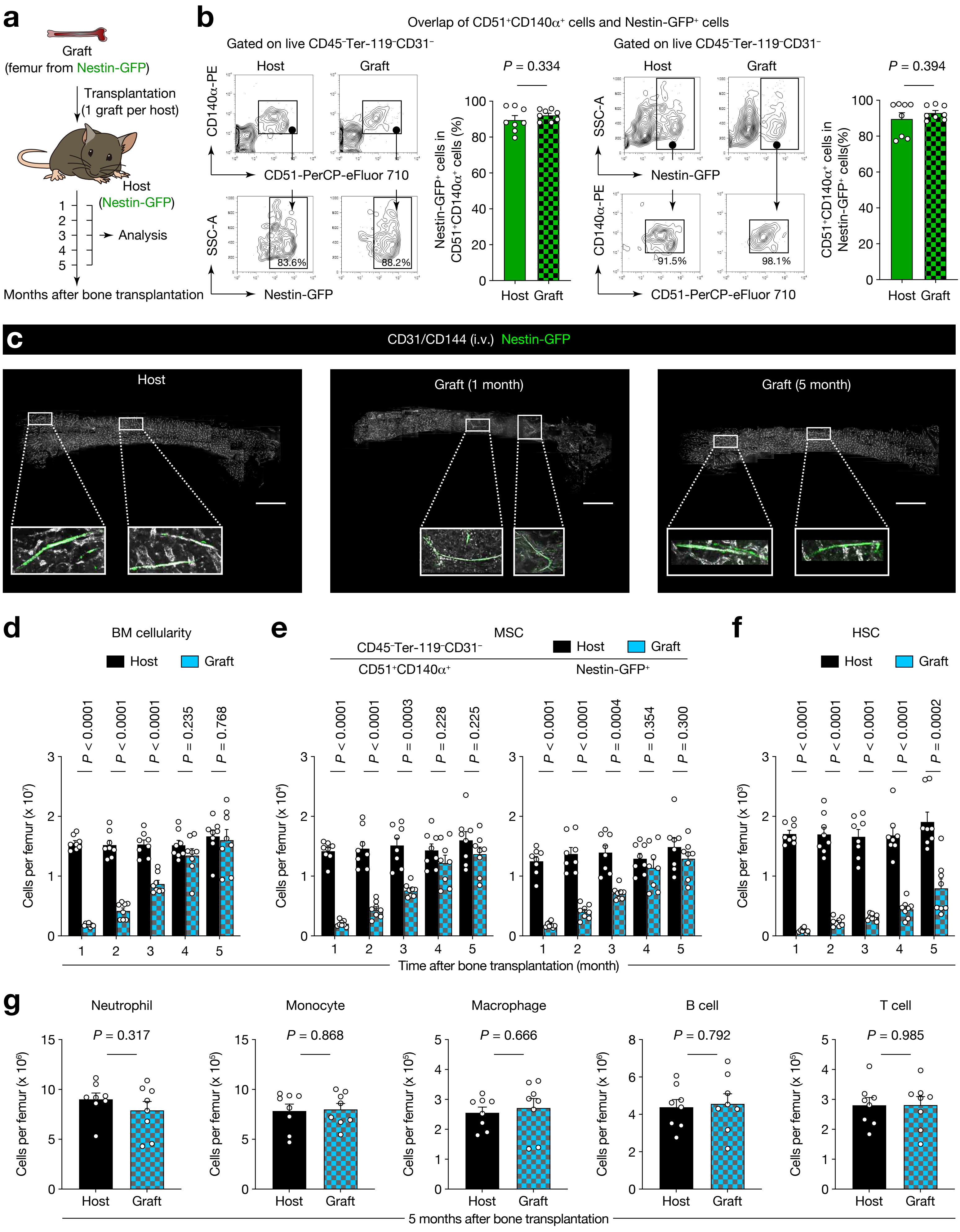


**Takeishi et al. Extended Data Fig. 4**

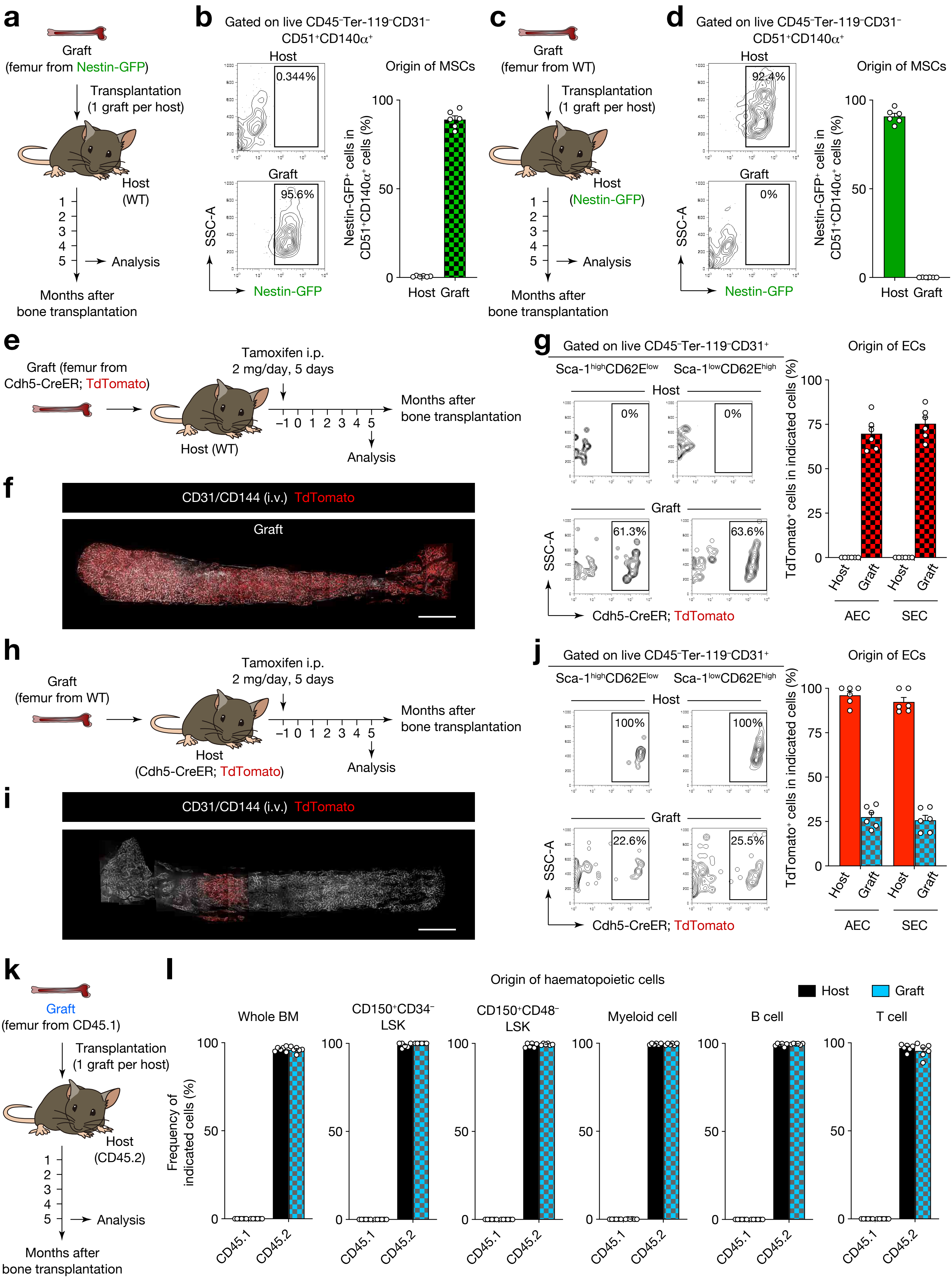


**Takeishi et al. Extended Data Fig. 5**

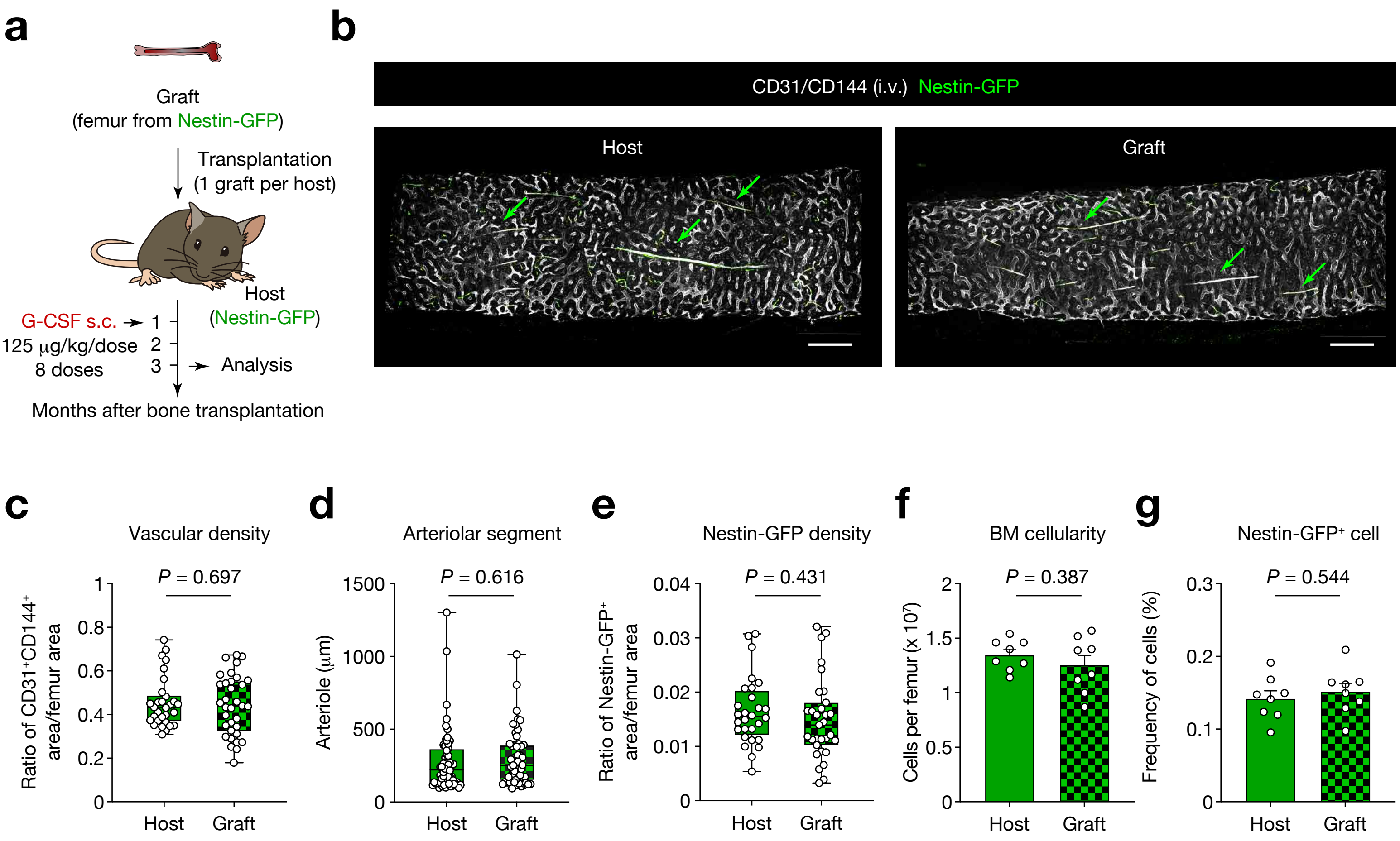


**Takeishi et al. Extended Data Fig. 6**

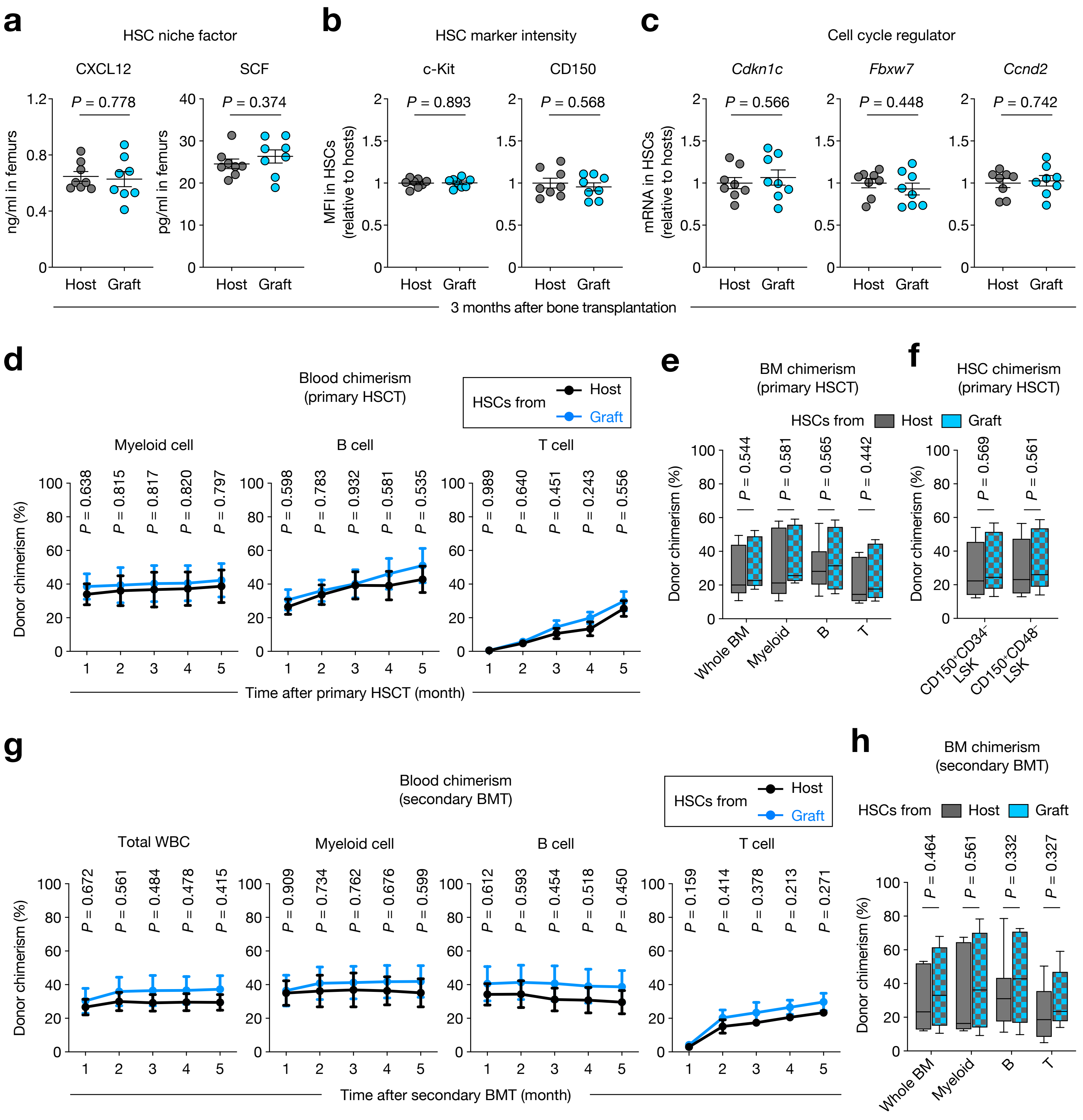


**Takeishi et al. Extended Data Fig. 7**

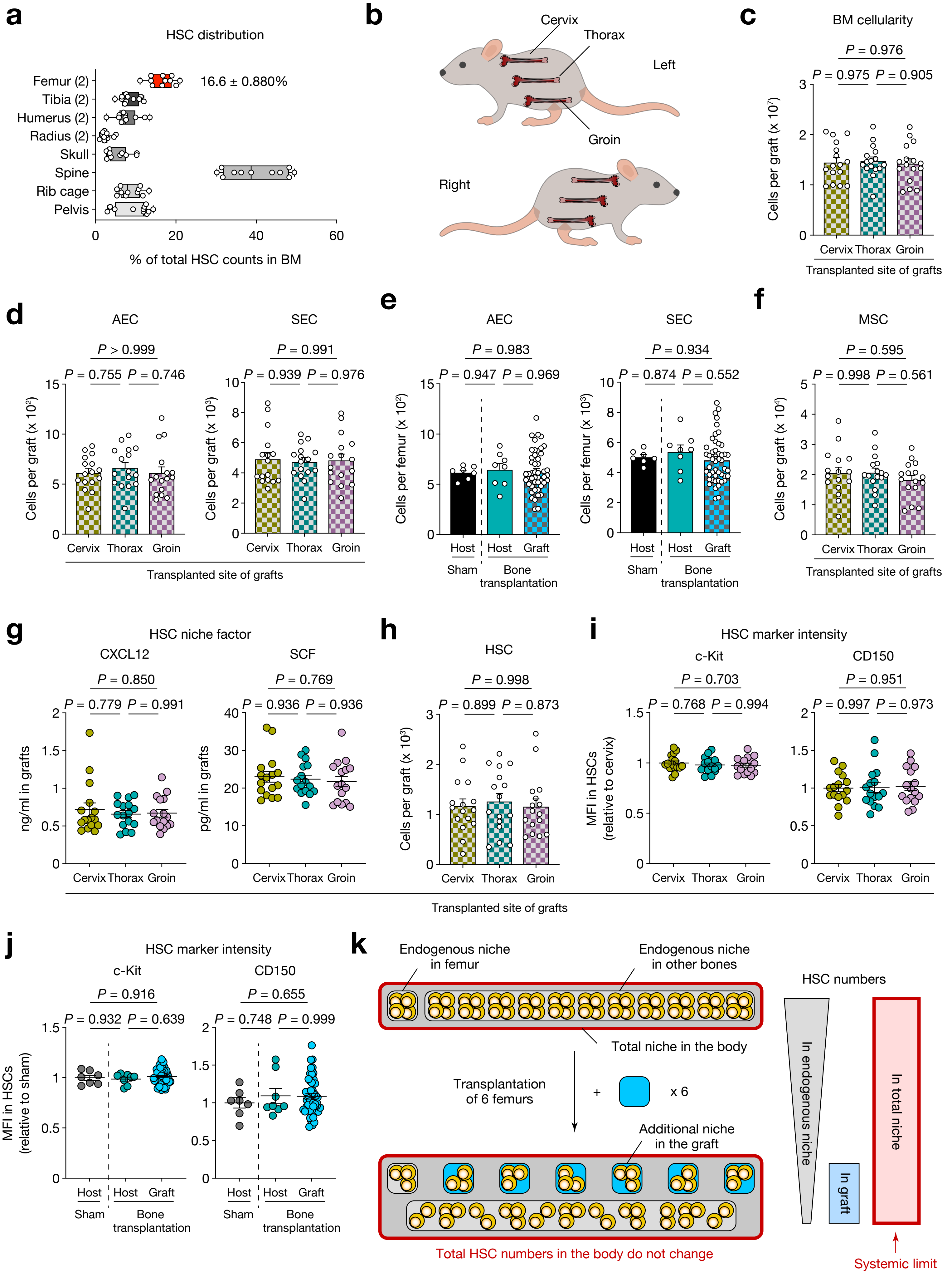


**Takeishi et al. Extended Data Fig. 8**

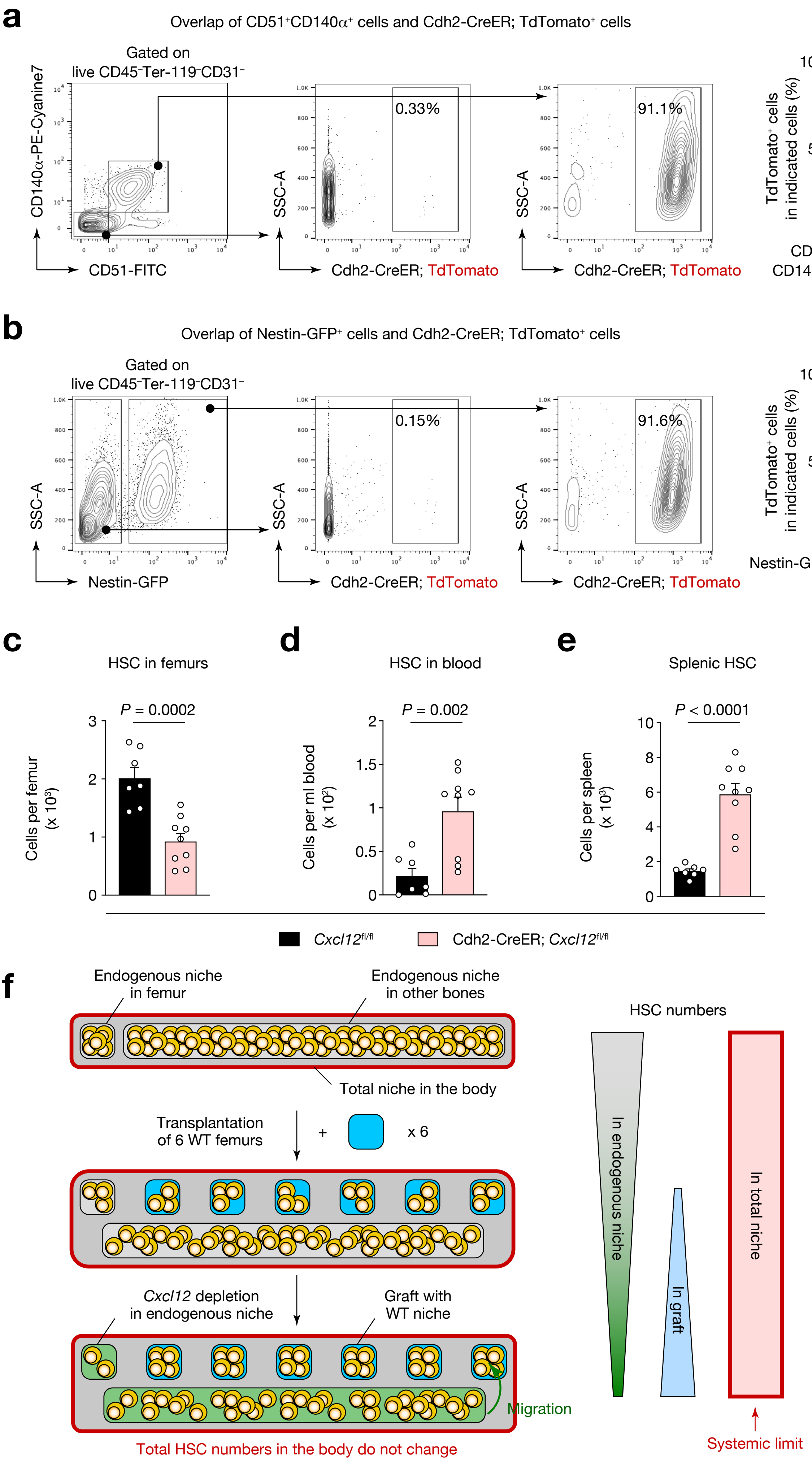


**Takeishi et al. Extended Data Fig. 9**

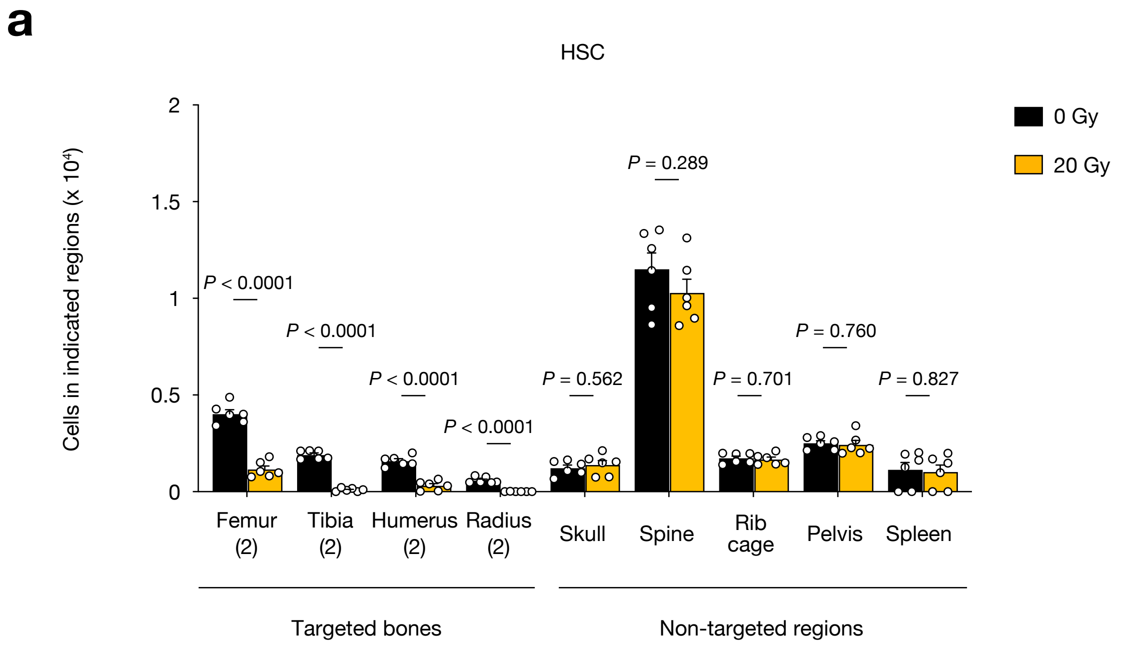


**Takeishi et al. Extended Data Fig. 10**

**a**

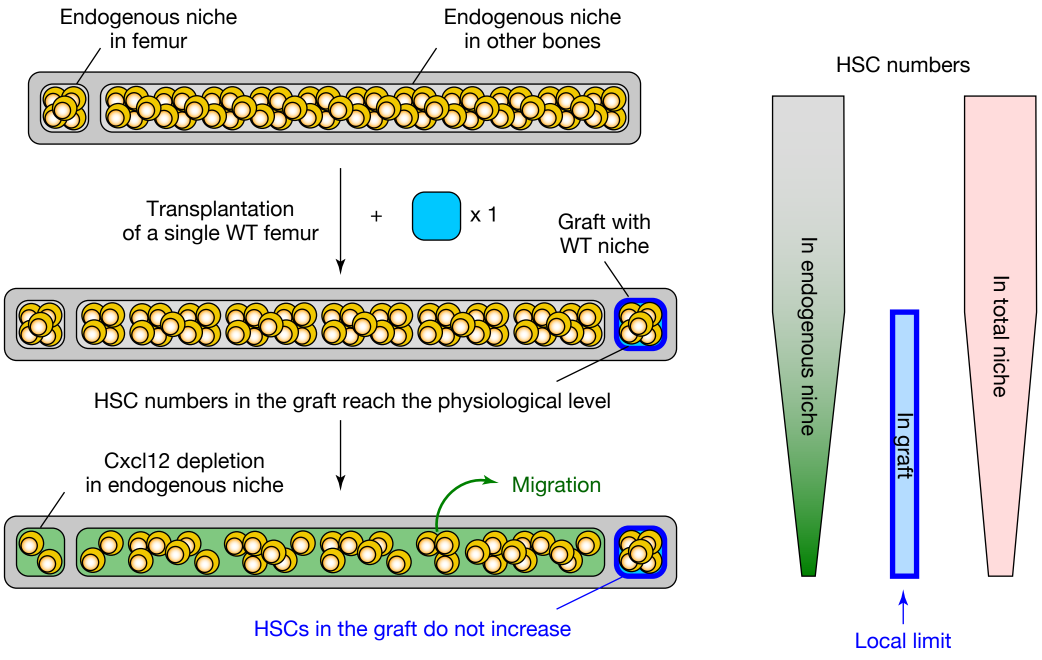


**Takeishi et al. Extended Data Fig. 11**

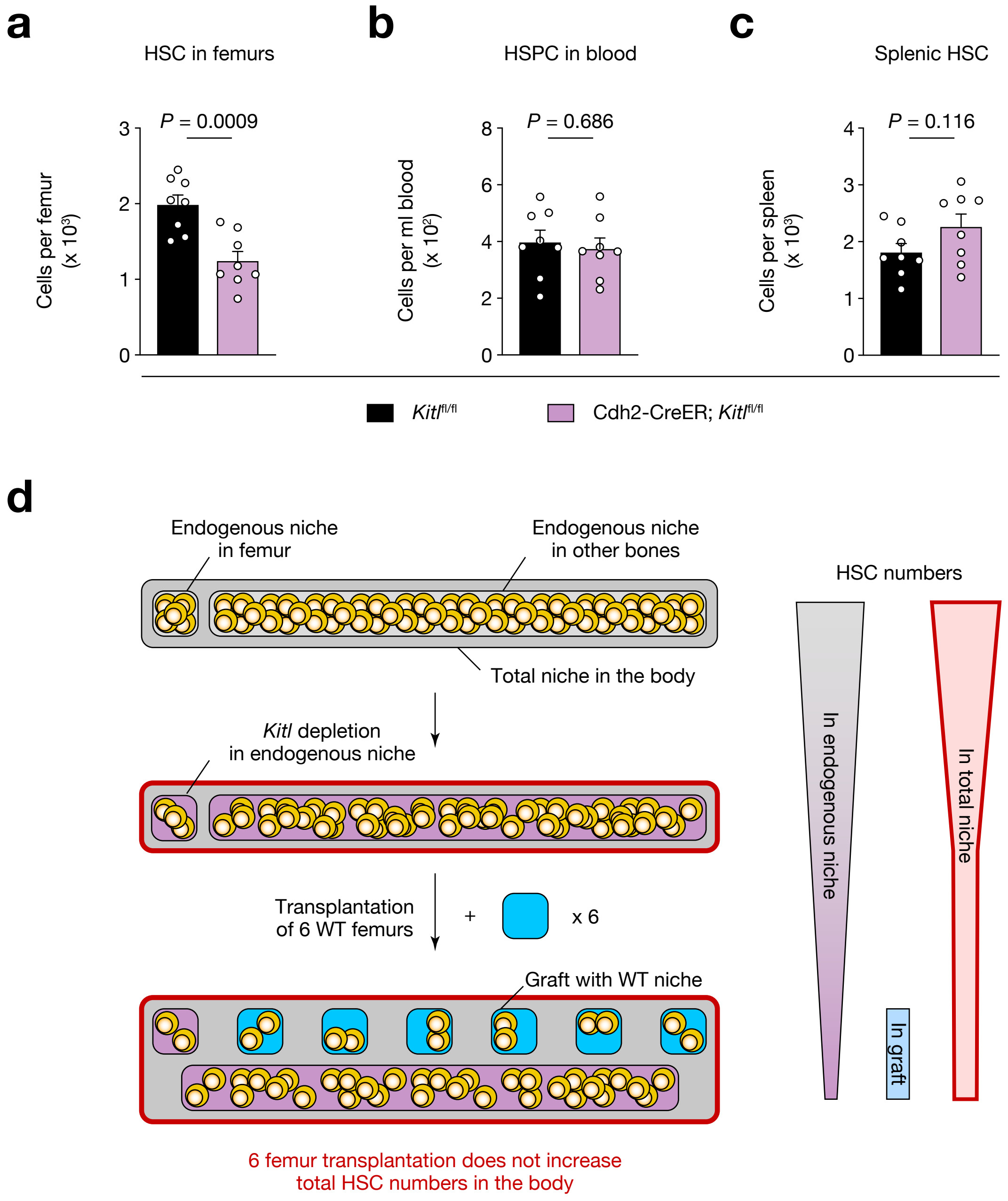


**Takeishi et al. Extended Data Fig. 12**

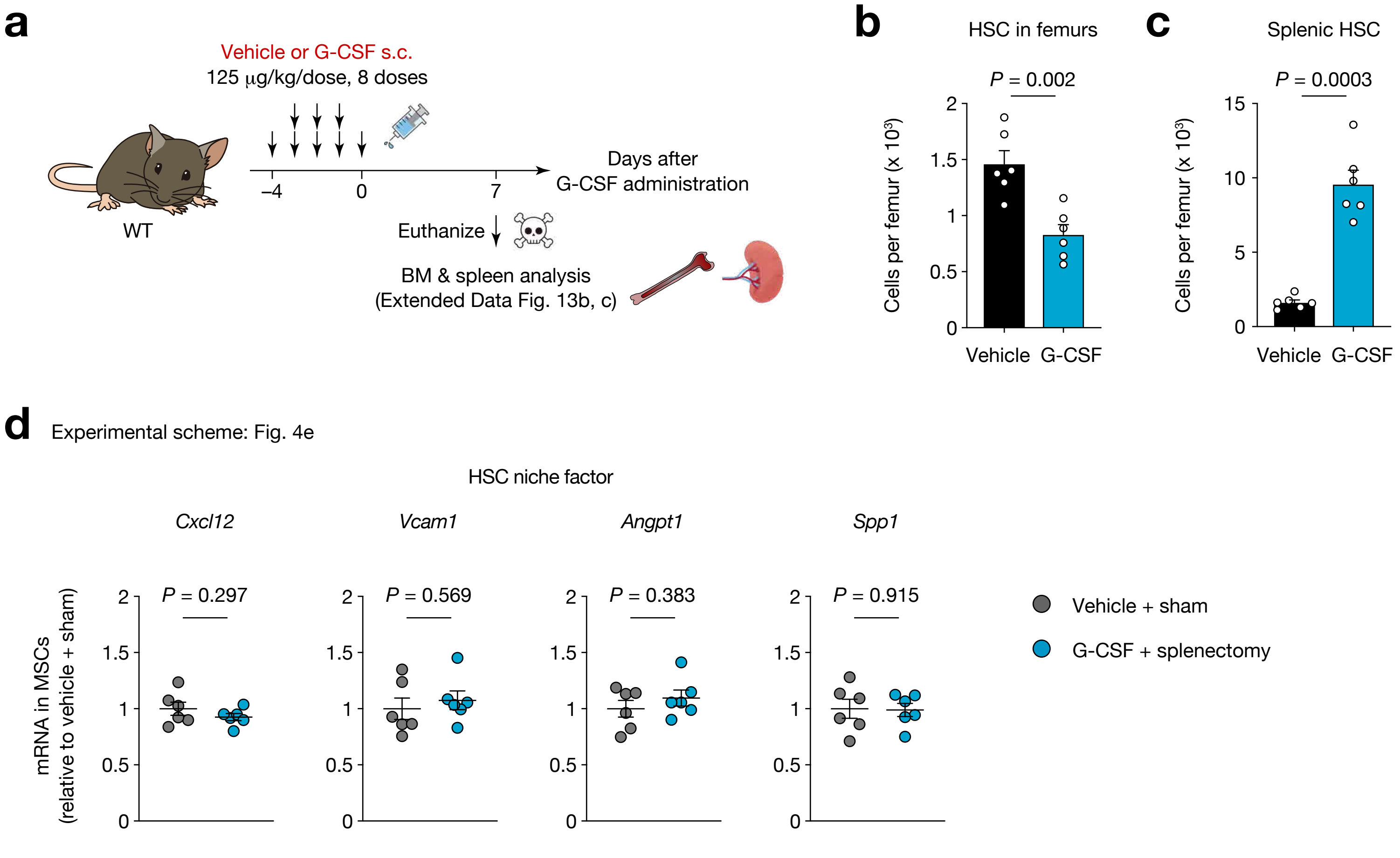


**Takeishi et al. Extended Data Fig. 13**

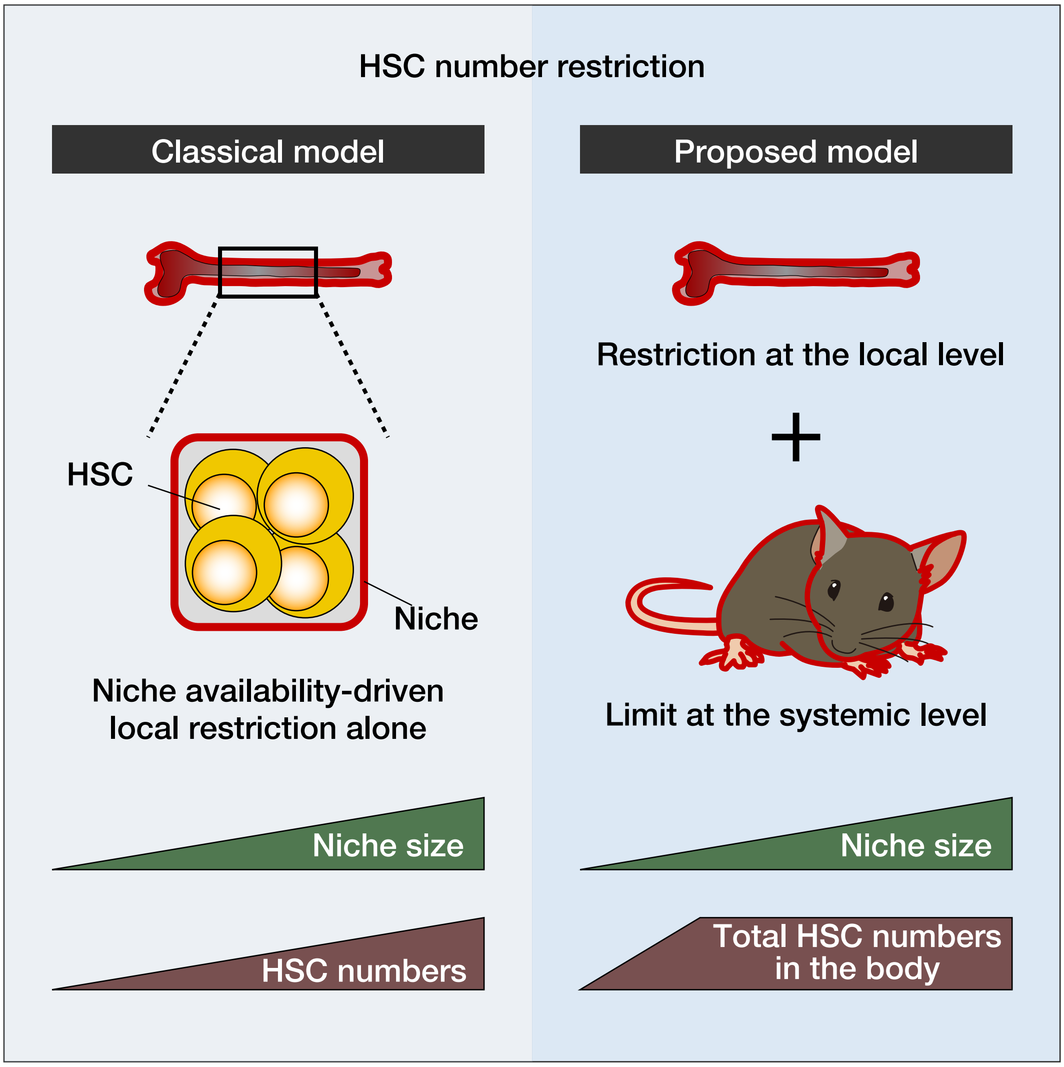


**Takeishi et al. Extended Data Fig. 14**
